# Activation of HIF2 leads to vascular remodeling and inflammation, coronary thrombosis and arterial dilation, recapitulating cardiac involvement of Kawasaki disease

**DOI:** 10.1101/2024.01.22.576642

**Authors:** B Escobar, I Menendez-Montes, T Albendea-Gomez, S Mendoza-Tamajón, R Castro-Mecinas, C Diaz-Diaz, B Palacios, J Ruiz-Cabello, LJ Jimenez-Borreguero, MC Cid, K Takahashi, S Martin-Puig

**Affiliations:** Cardiovascular Regeneration Program. National Center for Cardiovascular Research. 28029, Madrid, Spain; Department of Internal Medicine, University of Texas Southwestern Medical Center, Dallas, Texas, USA; Metabolism and Cell Signaling Department. Biomedical Research Institute Alberto Sols (IIBM). National Research Council (CSIC) – Autonoma University of Madrid (UAM). 28029 Madrid, Spain; Facultad de Medicina. Universidad Francisco de Vitoria, Madrid, Spain; Center for Cooperative Research in Biomaterials (CIC biomaGUNE), Basque Research and Technology Alliance (BRTA), 20014, Donostia, San Sebastián, Spain; CIBER de Enfermedades Respiratorias (CIBERES). Depart Química en C. Farmacéuticas. Faculty de Farmacia. Universidad Complutense de Madrid (UCM). 28040 Madrid, Spain; Cardiology Unit, Hospital Universitario de La Princesa, Madrid, Spain; CIBER Enfermedades Cardiovasculares (CIVERCV); Vasculitis Research Unit, Department of Autoimmune Diseases, Hospital Clinic, University of Barcelona, Institut d’Investigacions Biomèdiques August Pi i Sunyer (IDIBAPS), CRB-CELLEX, Barcelona, Spain; Department of Pathology, Toho University Ohashi Medical Center, Ohashi 2-17-6, Meguro-ku, Tokyo, 153-8515, Japan

## Abstract

**Background:** global deletion of *Vhl* leads to vascular defects and early lethality, precluding the study of VHL/HIF signaling during coronary formation and homeostasis. Hypoxia pathway has been associated with cardiovascular diseases involving inflammation and vascular remodeling like atherosclerosis, but its role in Kawasaki Disease (KD) remains unknown. Coronary dilatation and vessel rupture are the most serious complications of KD, while the molecular mechanisms underlying these cardiac events remain poorly understood. Here we aim to determine the function of VHL/HIF pathway in the development of cardiovascular defects and its role in KD.

**Methods:** We generated a new mouse model to genetically hyperactivate hypoxia pathway in progenitors contributing to coronary vessels and cardiac fibroblasts (*Vhl*/Wt1). We characterized the model by means of echocardiography, magnetic resonance imaging, histological analysis and molecular approaches. Human cardiac tissue from KD individuals suffering fatal coronary aneurysm were screened for HIF signaling and inflammatory markers by immunohistochemistry.

**Results:** conditional *Vhl* KO do not undergo developmental abnormalities but displays cardiomegaly and epicardial vascular defects, with cardiac hypertrophy and progressive coronary diameter increase, as well as pericardial hemorrhage and systemic inflammation early after birth. Histological characterization reveals inflammation of coronary arteries, vascular remodeling with elastin breaks and dilatation, increased perivascular fibrosis and smooth muscle cells death, together with high incidence of intracoronary thrombus formation. In addition, the mutants display vascular calcification and severe cardiac inflammation and interstitial hemorrhages, dying suddenly between 15-20 weeks of age due to vessel rupture. Simultaneous elimination of HIF2 and VHL prevents the cardiovascular abnormalities displayed by single c*Vhl* KO, highlighting the essential role of HIF2 in coronary instability and vascular inflammation. Histological characterization of human cardiac samples shows positive signal for HIF1 and specially HIF2, in the coronary lesions and its surroundings in regions with high inflammatory infiltration, confirming the activation of hypoxia signaling in KD patients with cardiovascular complications.

**Conclusions:** Our data demonstrate the importance of HIF2 signaling in the development of coronary inflammation and vascular remodeling and provide new evidences connecting low oxygen tensions with cardiovascular lesions occurring during the onset of the most severe cases of KD. Furthermore, the *Vhl*/Wt1 mouse generated recapitulates cardiac features of KD with critical heart complications, providing a new platform to uncover unknown aspects of KD pathogenesis.

## INTRODUCTION

Heart development is a complex process that rely on the participation of several progenitor pools that contribute to the diversity of the adult functional heart ^1,2^. Among them, epicardial progenitors represent a mixed population with potential to contribute to cardiac interstitial and adventitial fibroblasts, coronary smooth muscle cells and to a lesser extent to endothelial coronary cells ^3-6^. The potential of epicardial progenitors to give rise to the cardiac muscle lineage was controversial ^7^. Recent evidences indicate that the transcription factor Wilms tumor 1 (Wt1), a classical marker of epicardial progenitors ^5^, is also expressed in a subset of embryonic cardiomyocytes (CMs) that do not arise from epicardial-derived mesenchymal cells (EPDCs) during development ^8^. Within the heterogeneity of epicardial precursors ^9^, progenitors labelled by Wt1 represent one of the most broadly studied group contributing to the epicardium, interstitial fibroblasts, adventitial and medial layer of the coronary vessels ^5^, as well as to the leaflets of the atrioventricular valves ^10^. Interestingly, Wt1 expression has also been reported in endothelial cells (ECs) of cardiac capillaries in both mouse ^11^ and human ^12^ hearts.

Cardiogenesis occurs under relatively low oxygen levels or hypoxia at early stages, while as blood vessels develop during the formation of the coronary tree, the myocardium becomes better perfused and oxygen supply increases over time. HIFs are master transcription factors induced by hypoxia that in response to low oxygen concentrations mediate a broad transcriptional adaptive response, including enhanced expression of genes involved in glycolysis, erythropoiesis and angiogenesis among many others ^13^. In normal conditions the oxygen sensors prolyl hydroxylases (PHDs) mediate hydroxylation of specific proline residues within the alpha subunits of HIFs heterodimers. These posttranslational modifications serve as recognition marks for the HIF negative regulator Von Hippel Lindau (Vhl) that is part of a E3 ubiquitin ligase complex that facilitate HIF degradation through the proteasome after ubiquitination of the alpha subunits. In hypoxia PHDs become inactivated and, hence, HIF alpha subunits evade VHL recognition and further proteasome degradation, becoming stabilized and forming functional heterodimers with HIF beta subunits that translocate to the nuclei and binds to the promoter of target genes ^14^.

Mutations in *Vhl* have been associated with the development of different tumors, although our knowledge about the importance of VHL in cardiovascular physiology remains incompletely explored. In this regard, VHL has been reported to play a role in the regulation of vascular development. On one hand, *Vhl* global knock out mice display abnormal development of the placental vasculature and show embryonic lethality ^15^ and inactivation of *Vhl* by interference RNA causes rupture of the main embryonic vasculature and excessive looping of smaller branches ^16^. On the other hand, mutations in *Vhl* gene in the adulthood are associated with higher incidence of developing highly vascularized tumors like renal clear cell carcinomas, hemangioblastomas or pheochromocytomas ^17^. Endothelial-specific *Vhl*-null mice showed greatly dilated vessels with impaired vessel network patterning compared to control mice ^18^. Data from our laboratory indicates that full abrogation of *Vhl* in the myocardium impairs proper ventricular chamber development and cardiac maturation and results in cardiac dysfunction and embryonic lethality ^19^. These results point to the complexity of VHL function depending on the environmental, epigenetic and cellular context and suggest an important role for VHL in cardiovascular development and homeostasis. However, due to the early lethal phenotype of the former models, the role of VHL at later stages of cardiogenesis and the structural and functional consequences of *Vhl* deletion in epicardial progenitors contributing to coronary vasculature remains unexplored.

Kawasaki disease (KD) is a systemic vasculitis of medium size vessels affecting kids mainly from 0.5 months to 5 years old. It represents the main cause of acquired cardiovascular disease during childhood due to the high risk of coronary dilation and aneurysm development in untreated cases or in patients receiving delayed treatment ^20^. The combination of environmental and genetic factors has been proposed to participate in the progression of the pathology ^21^, but the etiology of this rare disease remains uncertain after over half a century of investigation since first described by Dr. Kawasaki ^22^. The development of animal models to investigate the pathogenic mechanisms of KD has contributed to identify the importance of innate immune response and IL-1β signaling in the development of vascular lesions ^23^. Currently, there are two available models based on systemic administration of L*actobacillus cassei* ^24^ or *Candida albicans* wall extracts ^25,26^. Nevertheless, the role of hypoxia and HIF signaling, which are important environmental factors, has not been evaluated in the development and progression of coronary vasculitis and KD.

Here we describe a novel mouse model of activation of hypoxia in epicardial progenitors and show that conditional elimination of *Vhl* in the Wt1 lineage has detrimental consequences in the integrity and stability of the coronary vasculature and cardiac homeostasis, leading to sustained inflammation, hypertrophy and fibrosis of the adult heart. Moreover, we discover that *Vhl/Wt1* mutants display stabilization of HIF1 and HIF2, coronary arterial inflammation, dilation and aneurysms, high rate of intracoronary thrombus formation and calcification associated with vessel rupture, describing a novel pathogenic role of HIFs in coronary vascular damage, and recapitulating cardiovascular defects typical of the most severe cases of KD. Moreover, we report the activation of HIF2 in KD cases with highly remodeled coronary vessels and damaged cardiac tissue, finding a positive correlation between hypoxia and vasculitis. Our findings contribute to uncovering the role of HIF signaling in the setting of this puzzling cardiovascular disease and the *Wt1/Vhl* deficient mouse model described here represents a valuable tool to explore molecular mechanisms underlying vascular inflammation and remodeling.

## MATERIALS AND METHODS

### Animal care and housing and human material

VHLflox/flox and HIF2 flox/flox mice ^27^ were obtained from Jackson laboratories (Strains 004081 and 008407 respectively) and maintained on the C57BL/6 background and crossed with mice carrying Wt1Cre recombinase ^28^ in heterozygosity. VHLflox/flox homozygous females were crossed with double heterozygous males and checked for plug formation. Mice were housed in SPF conditions at the CNIC Animal Facility. Welfare of animals used for experimental and other scientific purposes conformed to EU Directive 2010/63EU and Recommendation 2007/526/EC, enforced in Spanish law under Real Decreto 53/2013. Experiments with mice were approved by the CNIC Animal Experimentation Ethics Committee. Human cardiac sections used are from necropsies of Kawasaki disease cases from Toho Medical Centers in Japan and have been kindly provided by Dr. Kei Takahashi. Immunofluorescence and immunohistochemistry analysis performed on these tissues were approved by the Human Ethical Committee of the Toho University, Faculty of Medicine and 3 Medical Centers on April 2020.

### Genotyping

Genotyping was performed as previously described ^19^ using the following primers (Sigma Aldrich; USA) for VHL floxed alleles: Fw1: 5’ CTGGTACCCACGAAACTGTC 3’; Fw2: 5’ CTAGGCACCGAGCTTAGAGGTTTGCG 3’; Rv: 5’ CTGACTTCCACTGATGCTTGTCACAG 3’. For wt1 Cre alleles genotyping, the following primers were used: Fw: 5’TGA CG GTG GGA GAA TGT TAA T 3’; Rv: 5’ GCC GTA AAT CAA TCG ATG AGT 3’; For HIF2 floxed alleles: Fw: 5’ GAGAGCAGCTTCTCCTGGAA 3’; Rv: 5’ TGTAGGCAAGGAAACCAAGG 3’.

### Histological and immunohistochemical analysis

Histological sample processing and immunostaining was performed as previously described ^19^. Briefly, 5um thick paraffin sections were stained with hematoxylin & eosin (HE) following standard histological procedures at the CNIC Histopathology Facility. For immunostaining, sections were rehydrated, and antigens were retrieved by incubation in citrate buffer (10 mM sodium citrate, 0.05% Tween20, pH 6) in a pressure cooker. Sections were permeabilized with 0.5% TritonX100 for 10 min and blocked with 10% goat serum (GS) (Cat. No. 16210072, Life Technologies; NY; USA). Sections were incubated with primary antibodies in 10% GS overnight at 4°C. After several washes with PBST sections were incubated with secondary antibody (Life Technologies; NY; USA or Dako; Denmark) in 5% bovine serum albumin (BSA) for 1h at room temperature in the dark. When necessary, signal was amplified using the TSA System (Perkin Elmer; MA; USA). Finally, sections were incubated with DAPI (Millipore, MA; USA) and mounted in Fluorescent Mounting Medium (S3023, Dako, Denmark). Images were acquired with Zeiss LSM700 (Zeiss; Germany) or LEICA SP8 Navigator confocal microscopes. The primary antibodies used in this study were: HIF1α (NB100-479, Novus Biologicals; USA, GTX30647, Genetex, USA and CNIO (SIMA343B); Cy3-conjugated Smooth Muscle Actin (C6198, Sigma Aldrich; USA); EPAS (ab199 and ab109616, Abcam); ERG (ab199, Abcam), pH3 (06-670, Sigma-Aldrich); Ki67 (50-5698-82, Abcam), CD68 (MCA1957GA, BioRad), MMP9 (ab38898, Abcam), MRP14 (ab105472, Abcam).

### Quantification of histological sections and immunostaining

HE and Massońs trichrome staining was quantified as described previously) ^19^ using ImageJ ^33^.

### RNA extraction, cDNA synthesis and RT-qPCR

RNA extraction from hearts, cDNA synthesis and quantitative PCR were performed as previously described ^19^. Primers are available under request. Briefly, total RNA was extracted using QiAzol Lysis Reagent (Qiagen; CA; USA) and the miRNeasy Mini Kit (Qiagen; CA; USA). Total amount of isolated RNA was retrotranscribed using the MultiScribe Reverse Transcriptase kit 8 Applied Biosystems; CA; USA) and cDNA concentration was adjusted to 250ng/μL. All real-time qPCR reactions were performed in an AB7000 thermalcycler (Applied Biosystems; CA; USA) using SYBR Green PCR Master Mix (Applied Biosystems; CA; USA). Gene-specific primers were obtained from PrimerBank (http://pga.mgh.harvard.edu/primerbank/index.html) and checked for exon spanning using Primer3 (http://primer3.sourceforge.net/webif.php). Baseline normalization and thresholding were performed in automated mode with the SDS Software (Applied Biosystems; CA; USA). CT values were analyzed using qBase (Biogazelle; Belgium) using 3 housekeeping genes (HPRT and Rpl32). Primer-specific efficiencies were tested with serial dilutions of control cDNA. Statistically significant differences between control and mutants were analyzed by Student’s t test.

### Protein extraction and Western Blot

Tissues were homogenized using RIPA buffer and a TissueLyser in presence of protease and phosphatase inhibitors (Inhibitor cocktail (Roche, Switzerland) and 1μM sodium ortovanadate). After clarification by centrifugation, protein suspensions were sonicated and then concentration was measured using Pierce BA Protein Assay kit (23227, Thermo Scientific; USA) following the manufacturer instructions. 30μg of protein were denaturized at 95°C for 5 min, loaded on an 8% polyacrylamide SDS-PAGE gel and run at 120V for 90min. Subsequently, samples were transferred to a nitrocellulose membrane by wet transfer at 400mA for 2h. Membranes were blocked with 5% BSA for 1h and incubated with primary antibodies overnight at 4°C. Next day, membranes were washed in TBS-T buffer and incubated with the corresponding HRP-conjugated secondary antibodies (Dako, Denmark) at 1:5000 dilution for 1h at room temperature. After washing, signal was developed using ECL Primer Western Blotting Detection Reagent (Amersham; UK) and detected by a LAS-3000 imaging system (Fujifilm; USA). The primary antibodies used in this study were: anti-PAI-1 (sc-5297, Santa Cruz) 1:500 dilution and anti-vinculin (V4505, Sigma-Aldrich; USA) 1:5000 dilution

### Zymography

Heart extracts were homogenized using RIPA buffer and a TissueLyser in the presence of protease and phosphatase inhibitors (Inhibitor cocktail (11697498001, Roche, Switzerland) but in the absence of DTT. Cardiac extracts (30 µg) were resolved under nonreducing conditions on SDS–polyacrylamide gels containing 1% gelatin. Gels were washed three times in 2.5% Triton X-100 for 30 min at room temperature, incubated overnight at 37 °C in 50 mM Tris-HCl pH 7.5, 10 mM CaCl2 and 200 mM NaCl, and stained with Coomasie Blue. The areas of gelatinolytic or MMP activity were visualized as transparent bands. Images were analyzed with Quantity One software (Bio-Rad).

### Adult mice echocardiography and analysis

Mice were lightly anesthetized with 1 to 1.5% isoflurane in 100% oxygen and placed on a heated table to preserve physiological body temperature. Heart rate (HR) was monitored and anesthesia delivery was adjusted to maintain an HR of 500 ± 50 beats/min. Once anesthetic plane was reached, cardiac images were acquired for assessing cardiac dimension and function, by using a MS400 probe, at 30MHz for 2D and M mode images and 24MHz for Color and Pulsed Doppler modes, using an ultrasound scanner VEVO2100 (Visualsonics, Canada). Left ventricular ejection fraction (LVEF) by Teichholz formula was assessed using M-mode images displayed over time and obtained by a single line in the middle of LV. LV mass was calculated from the same M-mode images using diastolic LV diameters of LV internal diameter (LVIDd), LV posterior wall (LVPWd), and interventricular septum (IVSd) as follows: LV mass (mg) = 1.05 [(LVIDd + LVPWd + IVSd)3 − LVIDd3]. Corrected LV mass = (LV mass) 0.8 as previously described ^34^. To assess LV diastolic function, spectral pulse Doppler echocardiography was recorded for analyzing early (E wave) and late (A wave) velocities of mitral flow as previously described ^35^. To detect dilations of the proximal coronary arteries, images have been obtained at the level of the aorta root in axial and longitudinal planes using 2D echocardiogram and color Doppler.

### Magnetic Resonance Imaging

For cardiac MRI acquisition, animals were anaesthetized with isoflurane (2% pre and 1-1.5% during MRI acquisition) and their core body temperature, cardiac rhythm, and respiration rate were monitored using a MRI compatible monitoring system. In vivo cardiac images were acquired using an Agilent VNMRS DD1 7T MRI system (Santa Clara, California, USA). To obtain the images, a k-space segmented ECG-triggered cine gradient-echo sequence was utilized after shimming optimization. The protocol includes a four-chamber and two-chamber (long-axis view) of the left ventricle were captured and used to plan the short axis sequence. The mice were imaged using the following parameters: 13 slices, 0.8mm slice thickness with 0mm gap, matrix size of 256×256, field of view of 30×30mm2, ECG and respiratory triggered gating, 20 cardiac phases, 4 averages, ∼1.8ms minimum TE, 7ms minimum TR, 25^0^ flip angle, 2ms trigger delay, 8ms trigger window, and 2 dummy scans.

### Coronary resin cast filling and Computed Tomography analysis

Animals were deeply anesthetized (by 2% inhaled isoflurane) and i.p. heparinized, and then intravenously perfused with saline serum followed by injection of the MV122/133 microfil® suspension (Flow Tech Inc.) into the aorta until good filing of the coronary arteries is observed. This approach fills and opacifies microvascular spaces of forming a 3D cast of the vasculature filling with minimal shrinkage. After the solution had polymerized, the heart is removed, post-fixed in 4 % PFA overnight, and then stored in a 70 % alcoholic mixture (deyhol 70, 610070, Lavolan) at 4 ^0^C. CT image acquisition was carried out with a Nanoscan PET/CT (Mediso, Hungary) equipped with a microfocus X-ray source to generate 3D CT data of the heart with vascular casting, allowing for a comprehensive view and quantification of the heart vasculature. The images were acquired with 50 Kvp tube voltage and 145 µA beam current, 300 ms exposure time and 1:1 binning and 480 projections and reconstructed using proprietary Nucline software. The data was visualized and analyzed using Horos software (Horos Project, Switzerland).

### Statistical analysis and data representation

For histological, immunohistochemical quantifications, and RT-qPCR, values were pooled for mice with the same genotype and analyzed by the indicated statistical test using SPSS software (IBM; USA), with statistical significance assigned at P ≤0.05. Values were represented as mean±SEM using GraphPad Prism (GraphPad; USA).

## RESULTS

### Lack of Vhl in Wt1-coronary progenitors does not cause developmental abnormalities but leads to vascular defects, cardiomegaly and reduced survival early after birth

To investigate the importance of VHL protein in the establishment of coronary vessels during cardiogenesis, we generated a novel conditional mouse line by crossing the *Vhl*-floxed mice ^27^ with a *Wt1*-Cre line ^28^, from now on *Vhl* cKO mice. Histological characterization of *Vhl* cKO hearts by embryonic day E16.5 did not reveal any alteration in the coronary pattern or diameter, neither in chamber shape or width of the compact myocardium compared to controls (data not shown). Analysis of the offspring by weaning indicated that *Vhl cKO* mice survived embryogenesis and *Vhl-*deficient mice are recovered within the expected Mendelian ratios (Fig 1A). Efficient deletion of exon 1 of *Vhl* was confirmed by PCR (FigS1A), and resulted in reduction of *Vhl* mRNA levels and increased expression of the HIF target genes *Glut1, Phd3* and *Pai1* in hearts from *Vhl* cKO mutants compared to littermate controls (FigS1B). While the loss of one copy of *Vhl* gene in the Wt1 lineage did not result in any structural defect, neither in decreased viability (data not shown), the survival percentage rate was significantly compromised in the *Vhl* cKO homozygous mutants that displayed a high mortality rate with a median survival of 11.2 weeks of age (Fig 1B). Whole mount organ analysis revealed increased heart weight to body weight ratio (Fig1C), with evident cardiomegaly, right atria dilation and prominent vascular lesions present in the conditional *Vhl-*deficient mice (Fig1D, FigS2). These structural alterations did not result in cardiac systolic dysfunction and the *Vhl* cKO mice present normal ejection fraction (EF) (Fig1E) and fraction of shortening (FS) (Fig1F). In contrast, the early to late ventricular filling velocities ratio (E/A), indicative of diastolic dysfunction during cardiac relaxation, is significantly elevated by 12 weeks of age in the *Vhl* cKO mice (Fig1G), probably reflecting the increased interstitial fibrosis developed by the *Vhl* cKO (Fig3E). The gradual ventricular dilation, measured as left ventricular volume (Fig1H) and left ventricular inner diameter (data not shown) in the *Vhl* cKO mutants might be associated with the progressive cardiomegaly phenotype.

**Figure 1.**
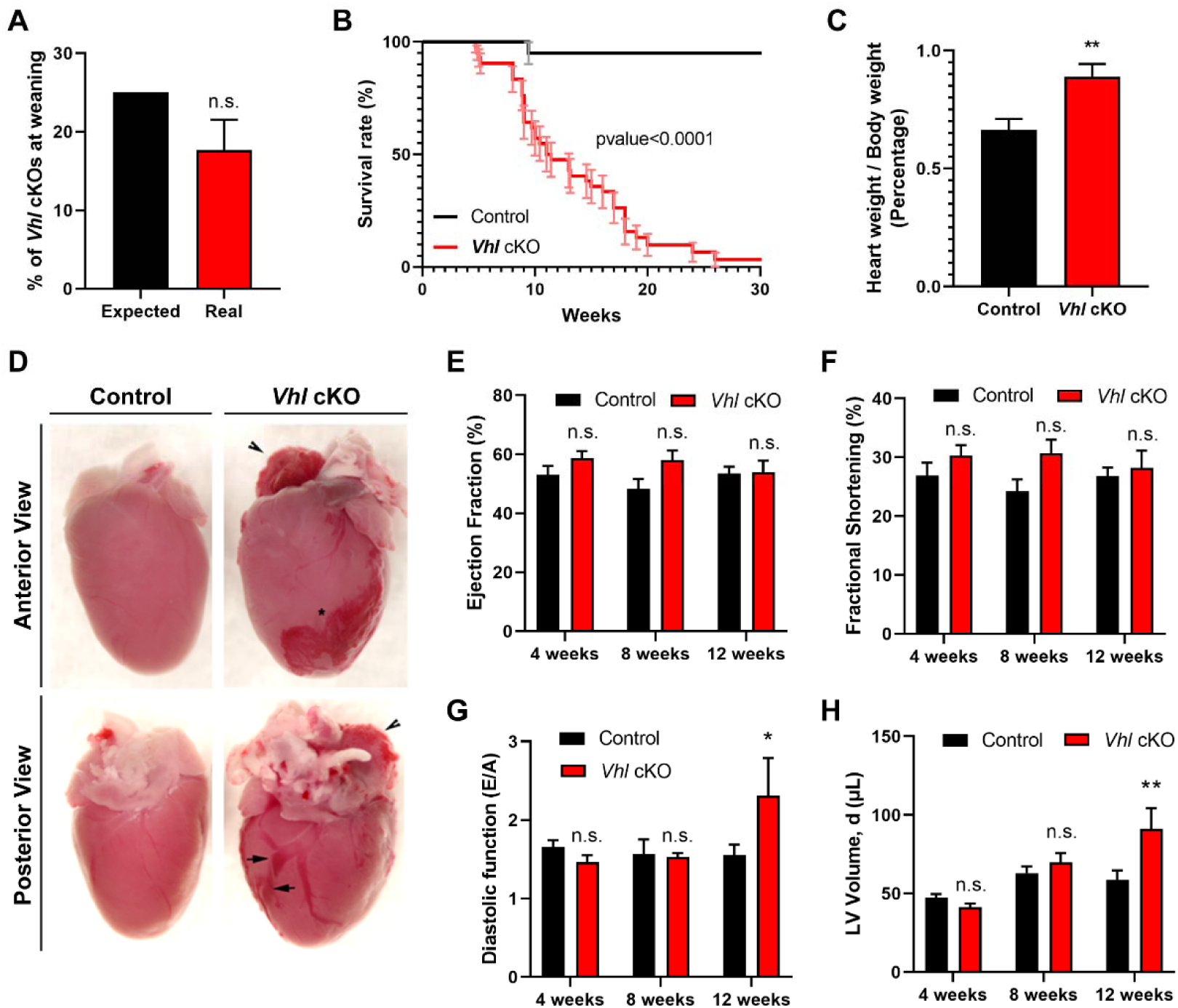
**Elimination of *Vhl* in the Wt1 lineage reduces adult survival, results in increased heart size and vascular lesions without reducing cardiac systolic function**. **A**). Percentage of recovery of *Vhl* cKO mutant mice by twenty-one days after birth. Quantification of the expected mendelian ratio based on the breeding strategy (black bar) and the real percentage of recovered *Vhl* cKO pups (red bar). Bars represent mean±SEM (n=27, real). n.s: non-significant. Statistical significance was tested by Student’s t test. **B**). Survival curve of control (black line) and *Vhl* cKO (red line) mice from birth to 30 weeks of age. n=73 (***P≤0.001). Statistical significance was tested by Log-rank (Mantel-Cox) text and Gehan-Breslow-Wilcoxon text. **C**). Heart size estimation based on the heart weight to body weight ratio of 8 weeks old mice. *Vhl* cKO mice (red bar) show cardiomegaly compared with controls (black bar). Bars represent mean±SEM (n=14 per group) (**P≤0,01). Statistical significance was tested by Mann Whitney test. **D**). Representative anterior (top panels) and posterior (bottom panels) whole mount views of hearts from control and *Vhl* cKO mice at 8 weeks of age. In the *Vhl* cKO arrowheads indicate right atria dilation, arrows show dilation of coronaries and asterisk indicates epicardial vascular lesions. **E-H)**. Longitudinal echocardiographic analysis of *Vhl* cKO (red bars) mice relative to controls (black bars) at 4, 8 and 12 weeks of age showing cardiac function measured by left ventricular ejection fraction (**E**), fractional of shortening (**F**) and early to late ventricular filling velocities ratio (E/A) (**G**). Dilation of the heart was estimated by measuring the left ventricular volume in diastole (**H**). All graphs represent mean±SEM. Control: 4 weeks (n=10), 8 weeks (n=7), 12 weeks (n=7). *Vhl* cKO: 4 weeks (n=10), 8 weeks (n=10), 12 weeks (n=9). Statistical significance was tested by Sidak’s multiple comparisons test (Two-way ANOVA).

In summary, these results indicate that epicardial *Vhl* is dispensable for proper coronary development during cardiogenesis, while its deletion results in acquired cardiac defects in the first weeks of life, leading to reduced adult survival, cardiomegaly and severe vascular lesions without affecting systolic cardiac function.

### Activation of HIF signaling triggers progressive coronary dilation and extracellular matrix remodeling

In addition to the evident vascular lesions of whole mount c*Vhl* KO hearts (Fig 1D), during the functional analysis by echocardiography we detected significant pericardial hemorrhage in the *Vhl* cKO mutants compared with controls (data not shown), suggestive of coronary defects in these mice. This finding was further observed by magnetic resonance imaging (MRI) that also confirmed enlarged cardiac size and revealed abnormally dilated coronaries in 8 weeks old *Vhl* cKO mice versus control hearts (Fig 2A). Further imaging analysis by echocardiography uncovered a clear enlargement of the coronary arteries in the *Vhl* cKO mutants without parallel alterations of the aorta (Fig2B). Longitudinal echocardiography analysis allowed quantification of the coronary diameter, showing a progressive increase in *Vhl* cKO mutants over controls between 4 and 12 weeks of age (Fig2C). Histological evaluation by hematoxylin and eosin staining corroborated that 8 weeks old *Vhl cKO* mutants displayed enlarged coronary arteries compared with controls, especially at the level of the proximal right coronary artery (RCA) (Fig2D). Interestingly, the *Vhl* cKO mutants did not display alterations in the medial layer of the aorta, which vascular smooth muscle cells (VSMCs) are not contributed by the Wt1 lineage (data not shown).

**Figure 2.**
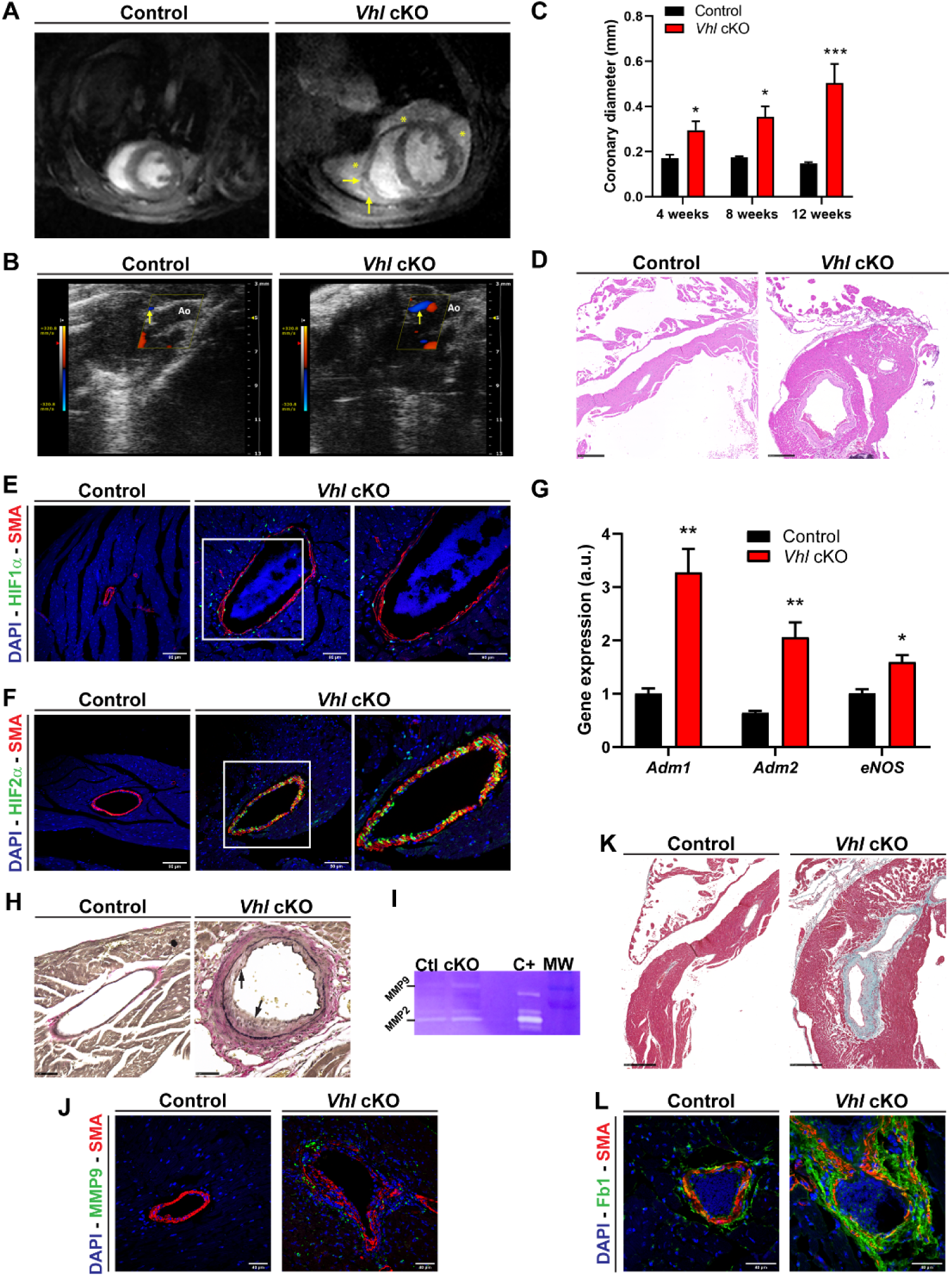
Activation of HIF signaling triggers progressive coronary dilation and enhanced extracellular matrix remodeling. A) Magnetic resonance imaging (MRI) show enlarged heart size of *Vhl* cKO mice with dilated coronaries (yellow arrows) and pericardial hemorrhage (yellow asterisks) relative to control. **B**) Echocardiography imaging revealed prominent enlargement of the coronary vessels (yellow arrows) in the *Vhl* cKO mutants without alterations of the aorta (Ao) compare with littermate controls. **C)** Longitudinal quantification from 4 to 12 weeks of age of the coronary diameter measured by echocardiography analysis showing a progressive enlargement in the *Vhl* cKO mutants (red bars) compare with controls (black bars). Bars represent mean±SEM. Control: 4 weeks (n=8), 8 weeks (n=5), 12 weeks (n=12). *Vhl* cKO: 4 weeks (n=8), 8 weeks (n=10), 12 weeks (n=8). *P≤0.05; ***P≤0.001; Statistical significance was tested by Two-way ANOVA test. **D)** Histological analysis of 8 weeks old adult cardiac sections by hematoxylin and eosin staining with zoom in the right coronary arteries of control and *Vhl* cKO mice. Scale bars 250μm. **E)** Immunofluorescence of cardiac sections against HIF1α (green), SMA (smooth muscle actin, red) and Dapi (blue) showing scarce HIF1α positive cells in fibroblast, ECs and VSMCs of the dilated coronaries in the *Vhl* cKO mutants compare with controls that stained negative for HIF1α. Scale bars 80μm. **F)** Immunofluorescence of cardiac sections against HIF2α (green), SMA (red) and Dapi (blue) reveals a strong signal of HIF2α labeling VSMCs and the endothelium and to lesser extent the adventitia of the *Vhl* cKO dilated coronaries compared with control. Scale bars 80μm. **G)** Relative gene expression analysis by RT-qPCR of the vasodilators *Adm1, Adm2* and *eNOS,* in control (black bars) and *Vhl* cKO mutant (red bars) heart lysates. Bars represent mean±SEM (n=3). *P≤0.05; **P≤0.01. Statistical significance was tested by Student’s t test. **H)** Representative 8 weeks old cardiac section analyzed by Verhoeff’s staining to define elastin (black lines) and collagen (pink) showing discontinuity in the elastic lamina (arrows) of the *Vhl* cKO mutant. Scale bars 50μm **I)** Representative zymography assay from 8 weeks old native cardiac lysates showing higher protein levels of matrix metalloprotease 9 (MMP9) and the active form of matrix metalloprotease 2 (MMP2) in *Vhl* cKO (lane 2, cKO) relative to control (lane 1, Ctl). C+ (lane 3) from an aortic lysate of a Marfan Syndrome mouse model; MW (lane 4) molecular weight. **J)** Representative immunofluorescence of MMP9 (green), SMA (red) and DAPI (blue) in 8 weeks old cardiac sections from control (left) and *Vhl* cKO mutants with expression of MMP9 between the adventitial and medial layer of a remodeled coronary (right). Scale bars 40μm. **K)** Representative 8 weeks old heart sections processed by Massońs trichrome staining in control and *Vhl* cKO mice showing increased collagen deposition (blue) in the enlarged coronary vessels. Scale bars 250μm. **L)** Immunofluorescence analysis of fibronectin 1 (Fb1, green), SMA (red) and DAPI (blue) in 8 weeks old cardiac sections reveals remodeling and perivascular fibrosis in the dilated coronary vessels of *Vhl* cKO mice relative to controls. Scale bars 40μm.

Since VHL is the negative regulator of HIFs, and the epicardial Wt1 lineage contributes to coronary endothelial and VSMCs, as well as to perivascular fibroblasts (FigS3A-D), we determined the expression of HIFs in the coronary arteries. Immunohistochemistry analysis showed scattered expression of HIF1α in ECs, VSMCs and fibroblasts of the dilated coronaries of the *Vhl* cKO mutants relative to controls (Fig 2E). Moreover, HIF2α levels were significantly increased in the coronary VSMCs, the endothelium, and to a lesser extent in the adventitia of the *Vhl* cKO mutant vessels (Fig 2F). In order to understand the molecular cues associated with these vascular abnormalities, we performed q-PCR analysis of endothelial nitric oxide (eNOS) that mediates the conversion of L-arginine into L-citruline and nitric oxide (NO), a potent vasodilator, finding that *Vhl* cKO displayed increased levels of eNOS relative to control. Furthermore, the expression of adrenomedullin (*Adm1*) and adrenomedullin 2 (*Adm2*), additional vasodilators modulated by hypoxia, was also elevated in the *Vhl* mutant compared to controls (Fig 2G). Next, we examined the pattern of elastic fibers in the coronary vessels of *Vhl* cKO mice by Verhoeff’s staining that revealed discontinuity in the elastic lamina of the *Vhl* cKO mutants, especially in regions of vascular hyperplasia and medial thickening (Fig 2H). To investigate pathways involved in vascular instability, we evaluated the activation of the matrix metalloproteases 2 (MMP2) and 9 (MMP9) that have proteolytic capacity of the extracellular matrix and have been previously described to be modulated by hypoxia ^29,30^. Zymography assay indicated an increase in both MMP9 protein levels and the active form of MMP2 in *Vhl* cKO mutant *versus* control hearts (Fig 2I). Furthermore, immunohistochemistry analysis showed that *Vhl* cKO mutant mice displayed increased expression of MMP9 between the adventitial and medial layer of the dilated coronaries compared with control vessels (Fig 2J). Moreover, we confirmed elevated levels of the tissue metalloproteinase inhibitor 1 (TIMP1) that is normally co-secreted as a complex with MMP9 (data not shown). To further investigate the extracellular matrix integrity of the coronary vasculature of *Vhl-*deficient mice, we performed Massońs trichrome staining of histological cardiac sections and observed prominent increased collagen deposition in the enlarged coronary vessels of *Vhl* cKO mutants compared to controls (Fig 2K), suggestive of perivascular fibrosis and remodeling. Moreover, immunohistochemistry against fibronectin 1 evidenced increased content in the dilated coronary vessels of *Vhl* cKO mutant relative to control hearts (Fig 2L).

In conclusion, these results demonstrate that loss of *Vhl* in Wt1^+^ epicardial progenitors results in progressive dilation of the coronary arteries associated with stabilization of HIF factors, increased levels of vasodilators like NO and adrenomedullins, as well as increased production of MMP9 and active MMP2, presence of elastin breaks and enhanced perivascular collagen and fibronectin deposition.

### Elimination of Vhl in the Wt1 lineage induces cardiac damage, capillary hemorrhages and fibrosis

Analysis of Wt1 reporter mice revealed positive signal in patches of CMs at the interventricular septum, papillary muscles and both ventricles (FigS3E-H), in addition to the contribution to microvasculature from these regions. The analysis of these structures by Masson trichrome staining in 4 weeks old mice reveals the initiation of tissue damage in CMs and the surrounding capillaries (FigS3I-L). Further histological characterization of 8 weeks old mice revealed foci of tissue remodeling and hemorrhagic dilated capillaries and interstitial fibrosis, especially in the papillary muscles and interventricular septum of *Vhl* cKO mutants relative to controls (Fig 3A). Furthermore, there is increased collagen deposition along ventricular and atrial epicardium in *Vhl* cKO mutants compared with controls (Fig 3B). This increased collagen content in *Vhl* cKO revealed interstitial fibrosis and correlated with the development of ventricular diastolic heart failure (Fig 1G). Indeed, longitudinal electrocardiographic analysis at different time points indicated significant prolonged PR interval and slight, but not significant, increased QRS segment by 12 weeks of age, (Fig 3C, 3D), suggestive of partially compromised atrioventricular and ventricular conduction over time respectively. These observations were consistent with the increased percentage of fibrotic areas in the *Vhl*-deficient mice (Fig 3E). Gene expression analysis by RT-PCR in 8 weeks old mice showed reduced expression of *elastin* together with increased levels of *fibronectin1*, *lisyl oxidase like 2* (Loxl2), *galectin 3* (Lgal3) and *transforming growth factor beta i* (Tgfbi), all involved in myofibroblasts activation (FigS4).

**Figure 3.**
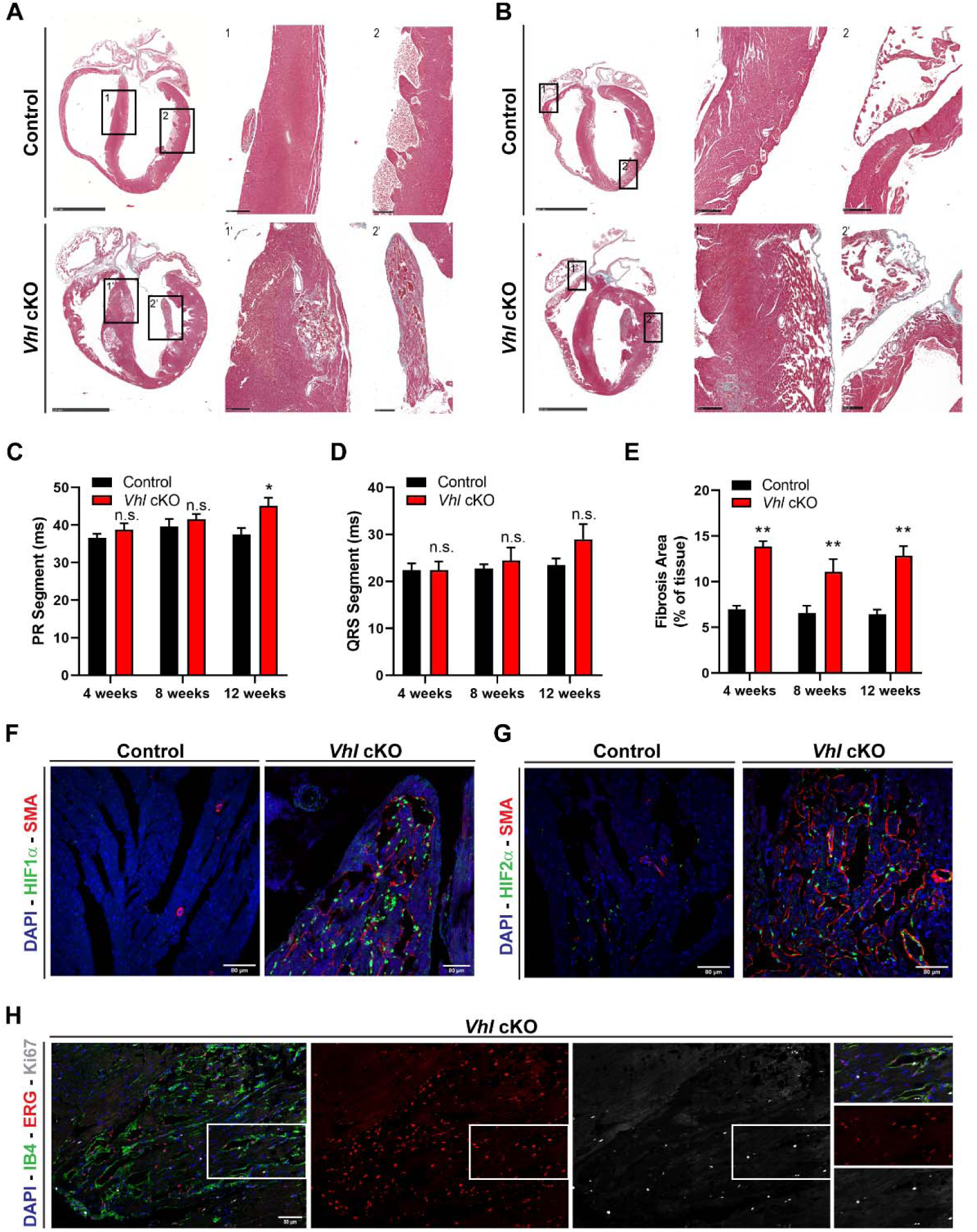
**Elimination of *Vhl* in the *Wt1* lineage results in cardiac damage, hemorrhagic capillaries and fibrosis. A-B**. Histological analysis by Massońs trichrome staining in 8 weeks old mice. **A)** Left panels show a representative overview of a cardiac section from control (top) and *Vhl* KO mutant mice (bottom). Scale bars 2,5mm. Middle (1, 1’) and right (2, 2’) panels show a magnified view of the interventricular septum (IVS) and papillary muscles respectively. Scale bars 250μm. **B)** Left panels show a representative overview of a cardiac section from control (top) and *Vhl* KO mutant mice (bottom). Scale bars 2,5mm. Middle (1, 1’) and right (2, 2’) panels show magnification of the ventricular and atrial epicardium respectively. Scale bars 250μm. **C-D)** Longitudinal electrocardiographic analysis of *Vhl* cKO (red bars) mice relative to controls (black bars) at 4, 8 and 12 weeks of age showing the PR (**C**) and QRS (**D**) Segments. Bars represent mean±SEM. Control: 4 weeks (n=10), 8 weeks (n=7), 12 weeks (n=7). *Vhl* cKO: 4 weeks (n=10), 8 weeks (n=10), 12 weeks (n=9). *P≤0.05; n.s: non-significant. Statistical significance was tested by Sidak’s multiple comparisons test (Two-way ANOVA). **E)** Quantification of the fibrosis area over time measured on cardiac sections of control (black bars) and *Vhl* cKO mutants (red bars) stained with Massońs trichrome. Bars represent mean±SEM. Control: 4 weeks (n=3), 8 weeks (n=7), 12 weeks (n=3). *Vhl* cKO: 4 weeks (n=5), 8 weeks (n=6), 12 weeks (n=4). **P≤0.01. Statistical significance was tested by Sidak’s multiple comparisons test (Two-way ANOVA). **F)** Immunofluorescence of HIF1α (green), SMA (red) and Dapi (blue) of 8 weeks old cardiac sections at the papillary muscles in controls (left) and *Vhl* cKO (right) mice. Scale bars 80μm. **G)** Immunofluorescence analysis of HIF2α (green), SMA (red) and Dapi (blue) of 8 weeks old cardiac sections at the papillary muscles in controls and *Vhl* cKO mice. Scale bars 80μm. **H)** Ki67 Immunofluorescence in 8 weeks old cardiac sections from *Vhl* cKO mice. ERG (ETS transcription factor, endothelial marker, red); Ki67, proliferation marker, white; IB4, isolectin B4, membrane endothelial marker, green and DAPI, blue. Scale bars 80μm.

Since elimination of *Vhl* should result in HIFs stabilization in the Wt1 lineage contributed structures, we explored the activation of HIFs by immunofluorescence in 8 weeks old hearts, finding that both HIF1α (Fig 3F) and HIF2α (Fig 3G) levels were significantly increased in the dilated capillaries of 8 weeks old *Vhl* cKO mutant hearts compared with controls. Furthermore, we detected proliferation of ECs of affected capillaries on the papillary muscle area (Fig 3H), suggesting that activation of HIFs in these microvessels surrounding myocardial tissue remodeling could promote the generation of new blood vessels through angiogenesis.

Altogether, these observations indicate that deletion of *Vhl* in the Wt1 lineage results in CM damage, microvasculature proliferation and enlargement of capillary vessels with increased collagen deposition and interstitial fibrosis, especially in the papillary muscles and interventricular septum, associated with HIF1α and HIF2α expression and EC proliferation. Moreover, elimination of *Vhl* leads to the induction of genes related to extracellular matrix remodeling and epicardial fibrosis.

### Stabilization of HIF signaling triggers cardiac inflammation and vascular damage

The histological characterization of *Vhl* cKO mutant hearts revealed the presence of inflammatory cells in the myocardium of *Vhl-*deficient mice, especially close to the fibrotic areas at the epicardium, surrounding the dilated and stenotic coronary vessels and altered papillary muscles (Fig 4A). Hence, we decided to explore the nature of the infiltrated cells and first performed immunofluorescence against macrophage and neutrophil markers using CD68 and MRP14 (S-100A9) antibodies respectively in combination with smooth muscle actin (SMA) to identify coronary vessels and fibrotic regions. The analysis of the stained sections revealed a profound infiltration of macrophages in areas with marked fibrosis in the epicardium, coronary arteries and papillary muscles (Fig 4B). In addition, significant clusters of neutrophils could be identified at similar locations compared with the lack of inflammation in control hearts (Fig 4C). Interestingly, dilated arteries with damaged medial layer displayed increased expression of myeloperoxidase (MPO), an enzyme stored in granules and released to the extracellular space during neutrophil degranulation, what could explain the vessel injury observed as a discontinuity on SMA staining in the affected coronaries (Fig 4D). Indeed, we could detect TUNEL positive signal, indicative of apoptosis, in the regions of discontinued SMA staining of the *Vhl* cKO mutant arteries, suggesting that the release of MPO by neutrophils could contribute to local medial layer injury and remodeling (Fig 4E). Analysis of the expression levels of several inflammatory mediators like interleukin 1beta (IL1β) and chemoattractant molecules like monocyte chemoattractant protein 1 (MCP-1/CCL2) or C-X-C Motif Chemokine Ligand 1 (CXCL1), implicated in neutrophil attraction, revealed an induction in *Vhl* cKO mice hearts (Fig 4F) consistently with the inflammatory infiltration found in these mutants.

**Figure 4.**
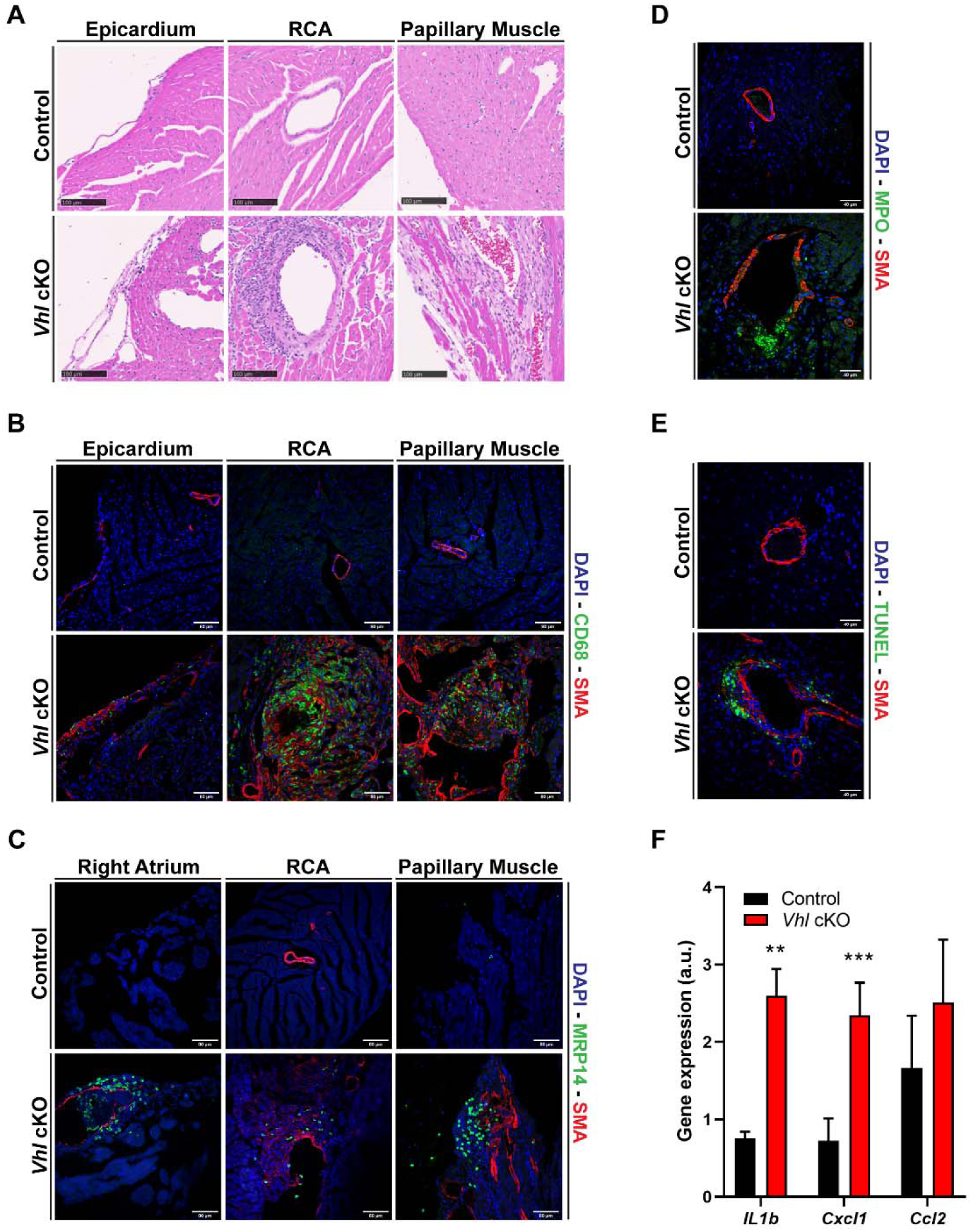
**Stabilization of HIF signaling promotes cardiac inflammation and vascular damage**. **A)** Histological analysis by hematoxylin and eosin staining in 8 weeks old control and *Vhl* cKO hearts. Representative views of epicardium (left), right coronary artery (RCA, middle) and papillary muscles (right). Scale bars 100μm. **B)** Immunofluorescence analysis of macrophages in 8 weeks old control and *Vhl* cKO hearts by immunostaining with CD68 (green), SMA (red) and DAPI (blue) in the same locations indicated in **A**. Scale bars 80μm. **C)** Immunofluorescence analysis of neutrophils labelled with MRP14 (green), SMA (red) and DAPI (blue) in 8 weeks old control and *Vhl* cKO mice at the same locations shown in **A** and **B**. Scale bars 80μm. **D)** Myeloperoxidase (MPO) protein expression around coronary vessels assessed by immunofluorescence of 8 weeks old cardiac sections from control and *Vhl* cKO mice. MPO (green), SMA (red), DAPI (blue). Scale bars 40μm. **E)** Apoptosis evaluation by TUNEL staining (green) in 8 weeks old cardiac sections from control and *Vhl* cKO mutants. Scale bars 40μm. **F)** Relative gene expression analysis of the inflammatory mediators interleukin 1 beta (*Il1b*), C-X-C Motif Chemokine Ligand 1 (*Cxcl1*) and chemoattractant molecule (*Ccl2*) by RT-qPCR from heart lysates of 8 weeks old control (black bars) and *Vhl* cKO (red bars) mice. Bars represent mean±SEM. Control: IL1b (n=2), Cxcl1 (n=7), Ccl2 (n=8). *VHL* cKO: IL1b (n=3), Cxcl1 (n=8), Ccl2 (n=9). **P≤0,01; ***P≤0,001. Statistical significance was tested by Student’s t test.

In summary, elimination of *Vhl* in the Wt1 lineage results in cardiac inflammation with prominent infiltration of macrophages and neutrophils that contribute to vascular damage and remodeling.

### Increased thrombus formation, vascular calcification and vessel rupture in Vhl-deficient coronaries

Further characterization revealed a high incidence of intracoronary thrombus formation in *Vhl* cKO mutant arteries compared with control hearts (Fig 5A). Thrombus are formed within cardiac cavities in several cardiovascular diseases like myocardial infarction or atrial fibrillation, but in mutants lacking *Vhl* the presence of thrombi was localized inside the coronary arteries, as observed in other human pathologies like atherosclerosis or KD. Moreover, in the biggest thrombi that fully occluded the lumen of the coronary vessel, we could observe the formation of nascent blood vessels, probably as a compensatory response to the compromised coronary perfusion upon occlusion, a phenomenon known as recanalization (Fig 5B) that also occurs in human pathological arteries. In addition, remarkable brown hemosiderin deposits are found in the thrombi, indicating hemolysis associated with vascular damage (Fig 5B, lower panels). Interestingly, the expression levels of plasminogen activator inhibitor 1 (PAI-1), a known target of HIF ^31,32^ involved in the fibrinolytic process, were significantly increased in the coronary vessels of *Vhl* cKO mutants relative to controls (Fig 5C, 5D), suggesting that a defective conversion of plasminogen into plasmin could contribute to the high incidence of coronary thrombus formation in the absence of functional VHL. Tridimensional evaluation of coronary vessels using a resin cast filling and subsequent analysis by computer tomography (CT) on 8 weeks old mice, revealed the appearance of aneurysm-like vascular lesions in *Vhl* cKO mutant hearts compared with controls (Fig 5E). This pattern progressed to uniform dilation or ectasia of the coronaries at later stages (data not shown). Finally, calcium deposits could be detected by Alizarin red or Von Kossa staining associated with dilated and remodeled arteries in the *Vhl* cKO hearts versus controls (Fig 5F and not shown), suggesting that enhanced rigidity and stiffness of mutant vessels could contribute to the coronary rupture, pericardial hemorrhages and sudden death observed in these animals.

**Figure 5.**
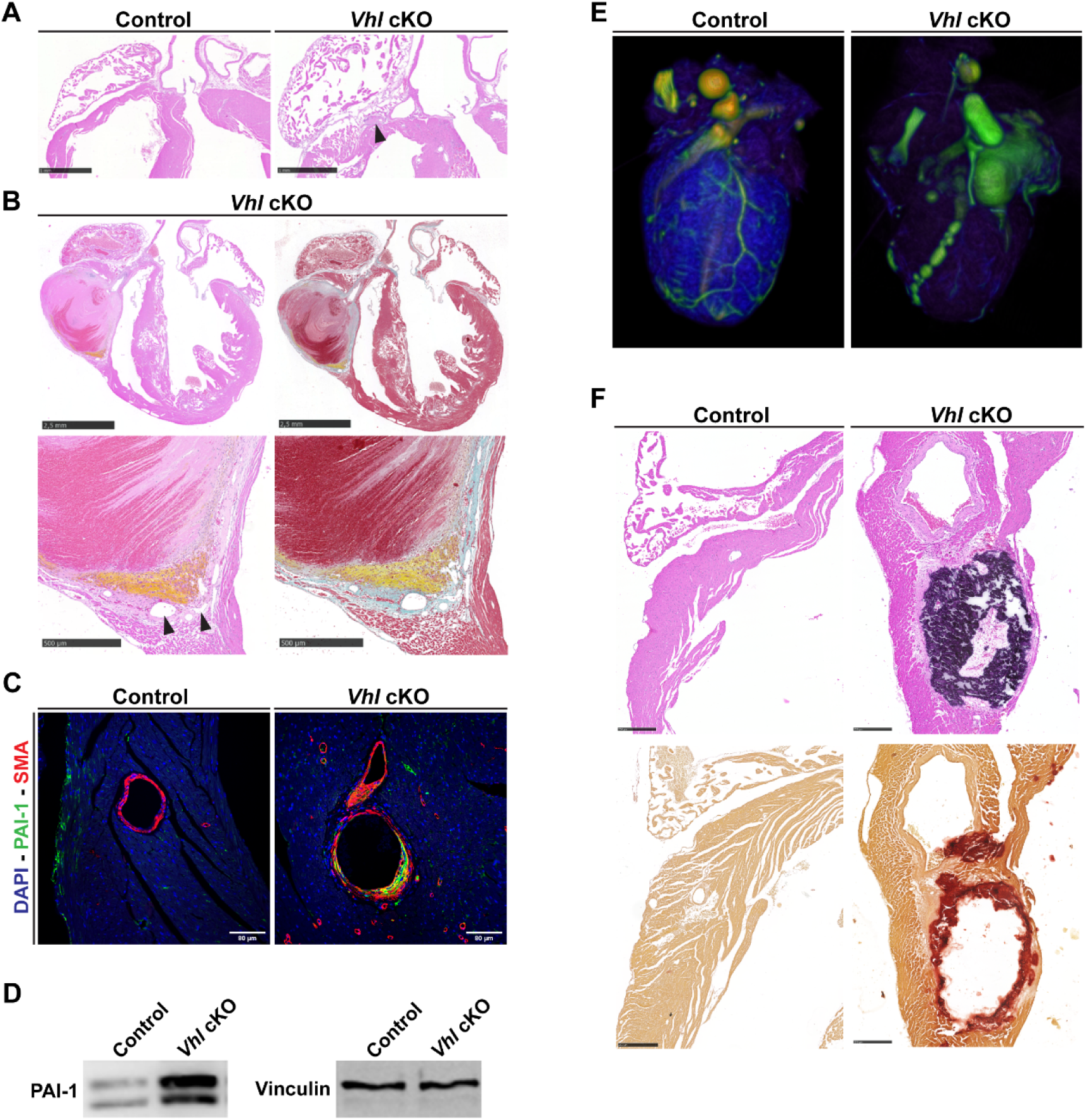
Increased thrombus formation, vascular calcification and vessel rupture in Vhl-deficient coronaries. A) Histological analysis by hematoxylin and eosin staining in heart sections from 8 weeks old control and *Vhl* cKO mice showing a magnified region of the right ventricle and atrium. The arrow in *Vhl* cKO mutant indicates the presence of a thrombus in the coronary artery. Scale bars 1mm. **B)** Histological analysis by hematoxylin and eosin (left panels) and Massońs trichrome (right panels) staining in heart sections of 8 weeks old *Vhl* cKO mice highlighting a large intracoronary thrombus near to the right coronary artery (top panels). Scale bars 2,5mm. Magnifications of the thrombus (bottom panels) show fibrin (pink) and collagen (blue) deposition, hemosiderin deposits (yellow) and nascent blood vessels (arrowheads). Scale bars 500μm. **C)** Immunofluorescence staining for plasminogen activator inhibitor-1 (PAI-1, green), SMA (red) and DAPI (blue) in coronary arteries from 8 weeks old control and *Vhl* cKO mice. Scale bars 80μm. **D)** Representative western blot of PAI-1 protein expression (left gel) from 8 weeks old control (lane 1) and *Vhl* cKO (lane 2) whole cardiac lysates. Vinculin was used as loading control (right gel). **E)** Coronary architecture evaluation by microfil resin-cast and computer tomography (CT) imaging analysis of 8 weeks old control and *Vhl* cKO mutant hearts showing aneurysm-like vascular lesions in the coronary arteries of *Vhl* cKO mice. **F)** Histological analysis by hematoxylin and eosin staining (top panels) in control and mutant (*Vhl* cKO) heart sections at the right coronary artery level. Alizarin red staining (bottom panels) to identify calcium deposits (red). Scale bars 250μm.

In conclusion, these observations demonstrate that the lack of *Vhl* in Wt1^+^ lineages induces the expression of PAI-1 and favors the formation of intracoronary thrombus, aneurysms and calcium deposits, replicating severe cardiac features typical of KD.

### Kawasaki disease patients display activation of HIFs around severe coronary vascular lesions

The analysis of our newly generated *Vhl* cKO mice indicates that activation of HIF signaling in the Wt1 lineage results in increased vascular remodeling, coronary dilation, aneurysm formation, intracoronary thrombosis, pericardial hemorrhage and myocardial inflammation and fibrosis. These cardiovascular defects are characteristic complications of the most severe cases of KD, a rare pediatric vasculitis of unknown origin and underexplored pathogenesis that represent the first cause of cardiovascular disease acquired during childhood ^21,22^.

Histological analysis of transverse sections of the ventricular wall of a representative necropsy from a 3 months old female case of KD shows a remarkable remodeling of the left anterior descending coronary artery (LAD) by hematoxylin and eosin staining, together with substantial immune infiltrate and intracoronary thrombus formation (Fig 6A, top inset), while remote left ventricular myocardium far apart from the damaged vascular lesion, does not display substantial abnormalities, neither immune infiltration or changes in capillary density, and conserves a healthy structure (Fig 6A, bottom inset). Immunohistochemistry analysis reveals significant nuclear expression of HIF2α in VSMCs and ECs of remodeled vessels and inflammatory cells around the vascular lesion (Fig 6B), as well as cytosolic HIF2α expression in arteries of the healthy myocardium (Fig 6D). In contrast, nuclear HIF1α was only detected to a fewer extent in some inflammatory cells close to the arterial lesion, but was barely expressed in VSMCs or ECs (Fig 6C) or around the healthy myocardium (Fig 6E). In another necropsy of a 7 months old male (Fig 6F) we confirmed the expression of HIF2 in VSMCs, ECs and inflammatory cells of an affected coronary (Fig 6F, top inset, and Fig 6G), as well as in macrophages surrounding the vascular lesion by immunohistochemistry with CD68 and CD163 (FigS5). In contrast, in agreement with the former female case, we could not detect expression of HIF1α in this lesion, neither in the inflammatory cells around it (Fig 6H). These observations were further confirmed in the proximity of a less remodeled artery in the area of healthy myocardium (Fig 6F, bottom inset), with substantial expression of HIF2α in VSMC, ECS and macrophages (Fig6I) without parallel detection of HIF1α (Fig6J). Similar HIF1α/HIF2α expression pattern profiles were observed in 3 additional cases of KD from 2 females and 1 male ranging from 3 months to 3 years old with fatal cardiac event after 10-30 days of the acute phase (data not shown).

**Figure 6.**
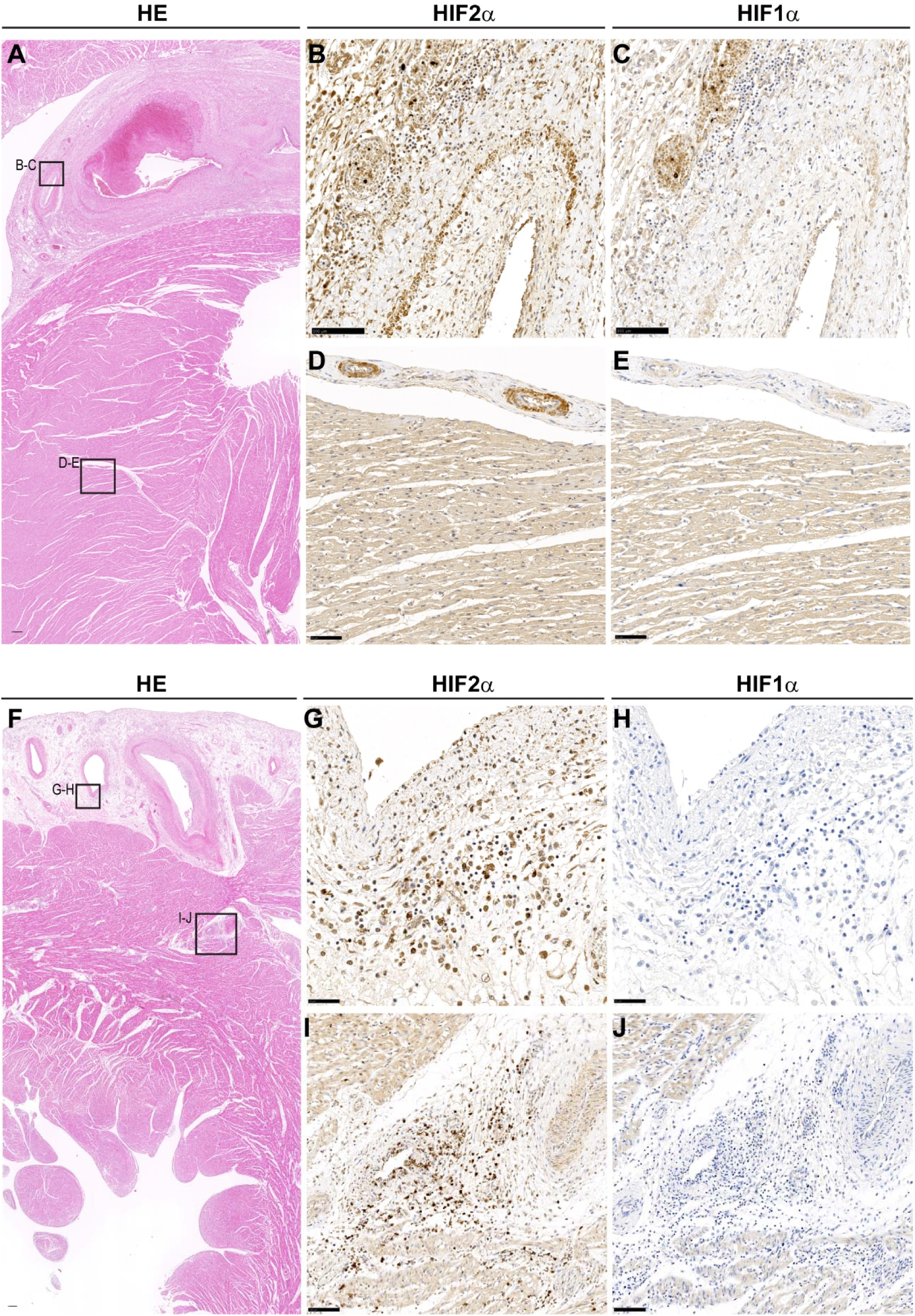
Activation of HIFs around severe coronary vascular lesions in Kawasaki disease patients. A) Histological analysis by hematoxylin and eosin staining of a representative heart section from a necropsy of a 3 months old female case of Kawasaki disease with vascular remodeling and thrombus formation in the descending left coronary artery (top inset, B-C) and healthy myocardium without signs of inflammation or vascular lesions (bottom inset D-E). Scale bar 250μm**. B-E**) Immunohistochemistry of HIF2α (**B**, **D**) and HIF1α (**C**, **E**) in the area of coronary remodeling (**B**, **C**. Scale bars 100μm) and in the healthy myocardium (**D**, **E**. Scale bars 50μm). Nuclei are counterstained with hematoxylin. **F)** Independent histological cardiac section from a necropsy of a 7 old months male case of Kawasaki disease stained with hematoxylin showing profound coronary remodeling without thrombus (top inset, G-H) and conserved left ventricular myocardium (bottom inset, I-J). Scale bar 250μm**. G-J)** Immunohistochemistry of HIF2α (**G**, **I**) and HIF1α (**H**, **J**) in the epicardial area of remarkable coronary remodeling (**G**, **H**. Scale bars 50μm) and around a partially-remodeled vessel in the ventricular myocardium (**I**, **J** Scale bars 100μm). Nuclei in **G**-**J** are counterstained with hematoxylin.

Collectively, these results indicate that the hypoxia pathway, specially HIF2 signaling, is active in the area of altered coronary vasculature in regions with exacerbated inflammation of patients with KD and might play an active role in the acute vasculitis and arterial remodeling of these patients, opening new avenues to the study of this unsolved pathology.

### Elimination of HIF2 rescues cardiomegaly, fibrosis, inflammation and coronary abnormalities of the Vhl/Wt1 mouse model

Because both the *Vhl* cKO mice and human cases of KD presented significant elevation of HIF2α in the area of coronary arterial lesions, we decided to generate a mouse model with double deletion of *Vhl* and *Hif2a* (*Wt1/Vhl/Hif2a mutant*) to evaluate the role of HIF2 in the abnormal hearts. *Wt1/Vhl/Hif2a* mutant mice –from now on, DKO-display a normal cardiac size and structure (Fig 7A), with a heart weight to body weight ratio comparable to control mice (Fig 7B). Echocardiographic studies revealed normal dimensions of all cardiac chambers (data not shown) with restored LV mass (Fig 7C) and coronary diameter (Fig 7D) on DKO compared with *Wt1/Vhl* single mutants. Histological analysis confirmed that the DKO mice have normal coronaries, without vascular remodeling or thrombus formation (Fig 7E), no alterations in the microvasculature of the papillary muscles (Fig 7F) and lack of atrial and epicardial fibrosis (Fig 7G) compared with *Vhl* cKO mutants. Furthermore, the extensive inflammation around the altered coronaries and the interventricular septum or papillary muscles observed in *Vhl* cKO mutant (Fig 4) is resolved in the absence of *Hif2a* (Fig 7H,7I).

**Figure 7.**
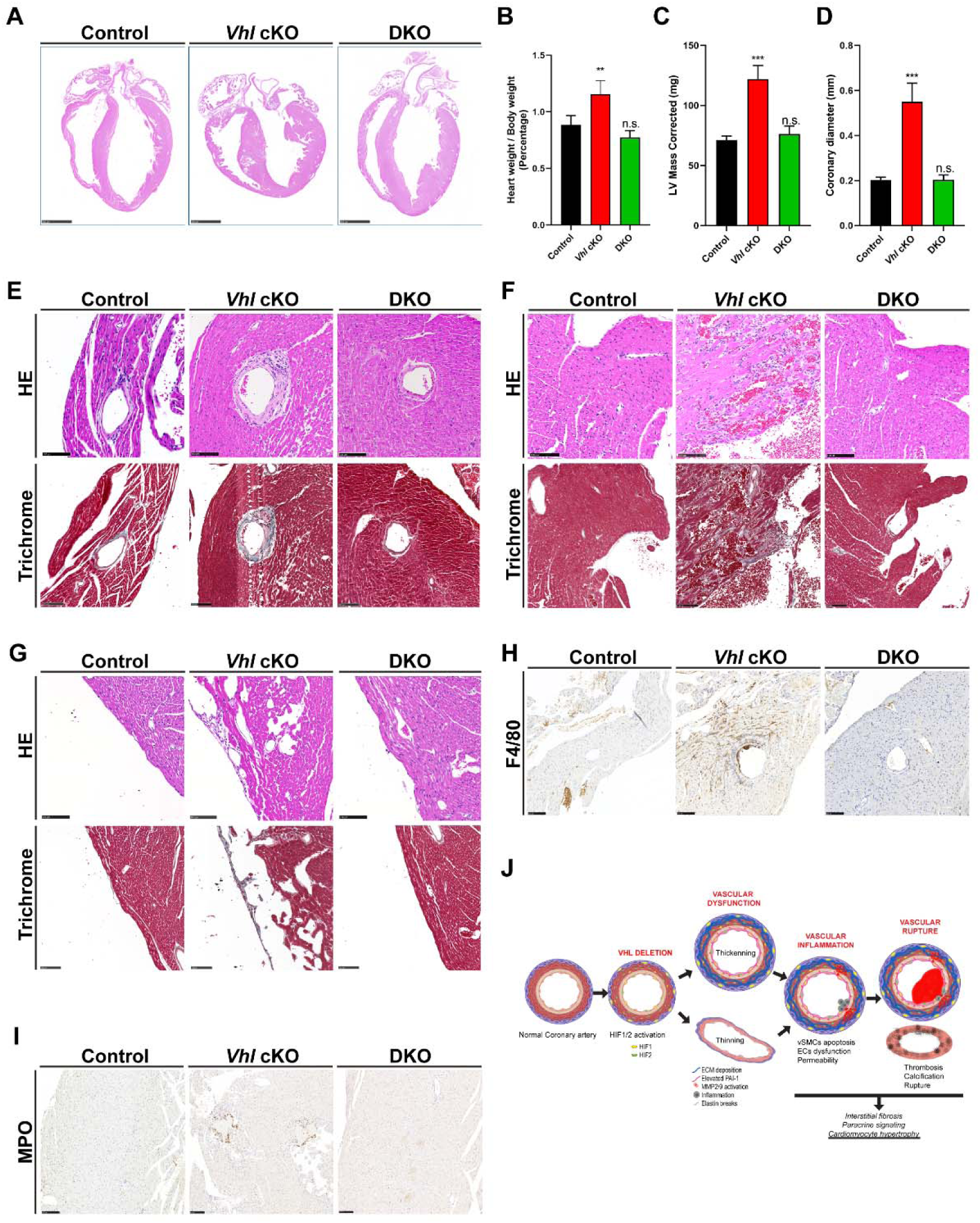
Elimination of HIF2 rescues cardiomegaly, fibrosis, inflammation and coronary abnormalities of the Vhl/Wt1 mouse model. A) Histological analysis of the cardiac structure in four chamber view by hematoxylin and eosin staining of control, *Vhl* cKO and double Vhl/Hif2a mutant (DKO). Scale bar 250μm**. B)** Cardiac size estimated as heart weight to body weight ratio of the indicated mice: control (black bars) (n=3), *Vhl* cKO (red bars) (n=8) and DKO (green bars) (n=3) mice. Bars represent mean±SEM. n.s: non-significant; **P≤0,01. Statistical significance was tested by Tukey’s multiple comparisons test (One-way ANOVA). **C-D)** Echocardiography analysis showing the left ventricular (LV) corrected mass (**C**) and the coronary diameter (**D**) in the indicated mice. Bars represent mean±SEM. Control: (n=4), *VHL* cKO (n=8), DKO (n=4). ***P≤0,001, n.s: non-significant. Statistical significance was tested by Tukey’s multiple comparisons test (One-way ANOVA). **E-G)** Histological analysis by hematoxylin/eosin (top panels) and Massońs trichrome staining (bottom panels) showing the right ventricular coronary artery structure (**E**), architecture of the interventricular septum and papillary muscles (**F**) and the epicardium integrity (**G**) in control, *Vhl* cKO and DKO mice. Scale bars 100μm. **H-I)** Representative immunohistochemistry of the inflammatory infiltrate by staining with F4/80 (**H**) and myeloperoxidase (MPO) (**I**) in the right ventricle of the indicated mice group. Scale bars 100μm. **J)** Proposed model of arterial vascular remodeling showing that deletion of Vhl in the Wt1 lineage leads to the activation of HIF1 and HIF2 in the coronary arteries. This genetic deletion promotes vascular dysfunction with increased extracellular matrix (ECM) deposition, elevated expression of Plasminogen Activator Inhibitor 1 (PAI-1) and Metalloproteases 2 and 9 (MMP2/9), causing elastin breaks and vascular remodeling (thickening & thinning of the arterial wall). These vascular alterations lead to VSMC apoptosis, ECs dysfunction and increased permeability and inflammatory infiltration. The *Vhl-*null arteries present a profound remodeling, with high incidence of intracoronary thrombus, calcification and tendency to rupture. In addition, the *Vhl* cKO mice display interstitial fibrosis and cardiac hypertrophy, most likely through indirect paracrine signaling.

Altogether these results demonstrate that constitutive activation of HIF2 signaling in cells of the Wt1 lineage upon deletion of *Vhl* causes vascular remodeling associated with increased activation of MMP2 and MMP9 protein, PAI-1 expression, abnormal ECM deposition and elastin breaks that results in vascular inflammation, VSMCs apoptosis, coronary and capillary dilation, vascular remodeling, which in turn favor the formation of thrombi and calcification of the arterial wall, eventually leading to vascular rupture, recapitulating cardiovascular complications of the most severe cases of KD (Fig 7J).

## DISCUSSION

Here we have explored the role of VHL and HIF signaling in the development and homeostasis of coronary vessels by characterizing novel genetic mouse models of the VHL/HIF axis in Wt1 progenitors. We demonstrate that elimination of *Vhl* in Wt1^+^ cells does not preclude normal coronary development but leads to severe vascular defects in young mice that present enlarged hearts and cardiac fibrosis, without affecting cardiac performance. Furthermore, *Vhl* cKO displays dilation of coronary arteries over time associated with HIFs stabilization, elevated expression of MMP2, MMP9, PAI-1, MPO and significant arterial remodeling and vascular damage, with increased collagen and fibronectin deposition, VSMCs loss and macrophage and neutrophil-mediated vasculitis. In addition, *Vhl* cKO displays high incidence of intracoronary thrombus formation and calcification, which together with high levels of collagen and fibronectin deposition around the inflamed vascular lesions, might increase rigidity and favor vessel rupture, contributing to the increased mortality of the mutant mice. Interestingly, the vascular and inflammatory defects upon HIF activation in the *Vhl* mutants recapitulate all the heart abnormalities of children with severe KD, a rare vasculitis representing the first cause of cardiovascular disease acquired during childhood. Here we uncover the expression of HIF2 in coronary VSMCs, ECs and inflammatory cells of the affected arteries in both male and female cases of fatal KD. Finally, we have demonstrated that HIF2 inactivation prevents vascular damage, inflammation and remodeling, rescuing all the cardiovascular abnormalities developed by the single *Vhl* cKO mouse model in the Wt1 lineage, pointing to the essential role of this HIF isoform in vascular stability and in the progression of cardiovascular conditions like KD.

### Cardiac hypertrophy, CM damage and fibrosis

Elimination of *Vhl* in the Wt1 lineage leads to cardiomegaly and CM hypertrophy. Considering that Wt1 is expressed in epicardial progenitors contributing to coronary VSMCs, some coronary ECs and cardiac fibroblasts (CF), as well as to capillaries and a subset of embryonic CMs, an important question is to determine whether the hypertrophy of CMs under *Vhl* elimination is a cell-autonomous or non-autonomous effect from surrounding cells of the microenvironment. On one hand, lineage tracing analysis of the Wt1 td-Tomato reporter mice reveals that the contribution of Wt1 to CMs is not homogeneous along the myocardium, but rather sparse, with higher contribution in the IVS and basal and apex areas of both ventricular chambers (SigS3E-H). Furthermore, our former model of *Vhl* deletion during cardiogenesis using Nkx2.5Cre as driver did not reach birth but did not result in increased size of CMs ^19^. Moreover, neither the lack of *Vhl* in neonatal CMs leads to an increased cell size (Diaz-Diaz C et al. unpublished observations), strongly suggesting that the cardiomegaly developed by the Vhl/Wt1 cKO model could be a non-cell-autonomous phenomenon. On the other hand, previous studies have demonstrated that angiogenesis could drive myocardial hypertrophy by secretion of VEGF-B or PR39 ^36^. Because *Vhl* deficiency in Wt1 lineage induces EC proliferation in capillaries, the secretion of paracrine factors from mitotic ECs to CM could be involved in the cardiac hypertrophy developed by the Vhl/Wt1 cKO. Indeed, it has been reported that several proangiogenic factors like nitric oxide (NO), neuregulin-1 (Nrg1), basic fibroblast growth factor 2 (bFGF2), platelet-derived growth factor (PDGF) and VEGF released from ECs contribute to cardiac hypertrophy and dysfunction ^37^. Furthermore, EC secreted factors such as endothelin 1, NO, Nrg1, Bmp4, Fgf23, Lgals3, Lcn2, Spp1 and Tgfb1 can induce cardiac hypertrophy and/or fibrosis ^38-41^. Interestingly, inducible deletion of the HIF-α upstream negative regulator PHD2 in ECs leads to left ventricular hypertrophy and fibrosis though a HIF2-dependent mechanism ^42^, suggesting that the activation of HIF2 signaling that we observed in the endothelium could be contributing to the cardiomegaly and cardiac hypertrophy as well as to the fibrotic phenotype of the Vhl/Wt1 cKO mice.

Another putative source of paracrine cues or cell-cell contact signaling contributing to the Vhl/Wt1 cKO phenotype could arise from CF derived from Wt1^+^ epicardial progenitors. Interstitial CF maintain the structural integrity of the heart through the regulation of extracellular matrix stability and remodeling. By modulating the levels of collagens, fibronectin and ECM remodeling factors like MMPs or TIMPs, CFs could directly impact on the microenvironment of CM. In response to cardiac damage CF undergo myofibroblast differentiation and adopt a pro-secretory phenotype that contribute to the release of proinflammatory and profibrotic cytokines and factors that might impact on cardiac hypertrophy and remodeling, including the conversion of CF into secretory myofibroblasts ^43^. Upon an ischemic insult, the increased expression and release of various cytokines, chemokines and growth factors such as IL6, IL1 beta, IL17, TGFβ, TNFα, MCP-1, endothelin-1, ANP, BNP, Ang II, FGF 2 or VEGF, among others ^44,45^ result in the modulation of both CFs and CM phenotype and promote inflammation ^43^. Thus, elevation of these factors, many of which are also HIF target genes, in CF could be involved in the profibrotic, proinflammatory and hypertrophic phenotype developed in the absence of Vhl in the Wt1 lineage.

In addition to cell-cell communication through paracrine cues like cytokines and growth factor, direct cell-cell contact and transcriptional regulation through microRNAs (miRNAs) released in exosomes could be involved in cardiac damage, fibrosis and hypertrophy observed in the Vhl/Wt1 cKO mice. In this regard, several miRNAs like miR-21, miR-155, miR-451, miR-146a and miR-200 have been reported to modulate cardiac hypertrophy (revised in ^46^). Defining the precise mechanisms underlying cardiac hypertrophy, fibrosis and inflammation in the Vhl/Wt1 cKO model will require additional analysis.

### Vascular permeability and instability/dilation

Regarding the vascular abnormalities developed by conditional elimination of *Vhl* in the Wt1 lineage, we have found that constitutive activation of HIFs in coronary arteries of Vhl/Wt1 cKO leads to vessel dilation as well as to profound vascular remodeling and inflammation. During exercise or low oxygen conditions coronary arteries dilate to favor blood flow increase. One of the postulated mechanisms of vascular dilation is by local accumulation of metabolic products like adenosine, lactate or acetylcholine ^47^. Since HIFs are master regulators of glycolysis and could mediate a metabolic reprograming in the coronaries, it is plausible that metabolic cues could be involved in the altered phenotype of the Vhl/Wt1 cKO. HIF also induces vasodilatation through increased expression of adrenomedullin, which is a vasodilatory peptide that stimulate endothelial NO release and hence mediates VSMCs relaxation ^48^. Because we observed increased expression levels of adrenomedullin 1 and 2, part of the dilated phenotype could be the result of these vasoactive mediators. The amount of NO has also been shown to be increased in cardiac ischemia in HIF-P4H-2 hypomorph mice through upregulation of endothelial nitric oxide synthase (eNOS) ^49^. In addition, eNOS expression is controlled by HIF2 ^50^ and is induced in the Vhl/Wt1 cKO (Fig 2G). Hence, elevation of NO levels could be involved in the coronary vasodilation observed in the mutants. Coronary dilation could also be associated with enhanced levels of MMP9/TIMP1 that might account for the elastin breaks of the remodeled coronaries observed in the Vhl/Wt1 cKO. Interestingly, MMP9 has been studied and implicated in elastin breakdown in the *Lactobacillus casei* cell wall extract (LCWE)-induced KD mouse model ^51^ and MMP9 and TIMP1 have been also found to be elevated in plasma of acute phase KD patients ^52^. Inducible elimination of Phd2 in adult mice leads to hyperactive angiogenesis and increased capillary density in the heart with enlarged blood vessels. Phd2-cKO mice also presents angiectasia in lung, liver and kidney ^53^. The enlarged vascular phenotype was observed in both veins and arteries in the Phd2 cKO model and only HIF1 but not HIF2 alpha levels were increased in the mutant relative to controls ^53^. Although systemic inactivation of Phd2 promotes angiectasia and angiogenesis, the phenotype is different from the Vhl/Wt1 cKO presented here that only display arterial but not venous dilation. Some of the differences between the Phd2 cKO and our tissue/lineage specific Vhl cKO might reflect distinct responses to systemic versus localized induction of HIF signaling in each mouse model and suggest that heterogeneous and local activation of HIF2 might contribute to the development of coronary arterial defects.

### Pathogenesis of KD

KD is a pediatric vasculitis of unknown etiology mostly affecting children between 0 and 5 years old that represent the main cause of acquired cardiovascular disease in childhood. The more severe cases develop coronary dilation and aneurysms, usually due to delayed treatment. In addition, a fraction of treated kids is refractory to immunoglobulin infusion being at higher risk of developing coronary lesions. The lack of specific diagnostic tools and treatments represent a challenge to reduce cardiovascular risk associated to late diagnosis or treatment. A better understanding of the molecular pathways involved in the pathogenesis of KD would highly contribute to the development of novel diagnostic and therapeutic strategies to minimize the complications and irreversible consequences of this devastating disease. Because there is a limited access to human damaged tissues from KD affected patients, the development and characterization of experimental mouse models of KD vasculitis has considerably improved our understanding of the pathology and helped to identify the cellular and molecular immune mechanisms contributing to its characteristic cardiovascular abnormalities ^26^. These studies have allowed to propose novel experimental treatments that are under evaluation in several clinical trials ^26,54-56^.

The current animal models used to study the molecular basis of KD are based on systemic administration of proinflammatory stimuli like LPS and either bacterial or fungal cell wall extracts, hence promoting a global inflammatory response not only in the coronary arteries. Because it has been described that KD patients with severe cardiac complications do not suffer aortic dilation despite of profound aortitis ^57^ the use of mouse models with global inflammation and systemic dilation of the vasculature might not fully reflect the clinical scenario of cardiovascular abnormalities encountered by some KD patients. In this regard, the novel Vhl/Wt1 cKO mice that we generated and characterized here recapitulates all the cardiac abnormalities existing in the most severe cases of KD, and to our knowledge represents the first genetic mouse model closely mimicking the cardiac alterations of this condition. The discovery of elevated HIF2 levels in the dilated arteries, as well as inflammatory cells surrounding the coronary lesions of KD cases, contributes to clarify the molecular pathogenesis involved in the progression of the disease. HIF signaling have been reported to modulate the immune response and could control phagocytic capacity, infiltration and chemotaxis. Therefore, our findings of elevated HIF2 levels in macrophages in the environment of KD damaged coronary arteries open novel research avenues towards our understanding of this puzzling disease.

## ARTICLE INFORMATION

## Acknowledgements

The authors thank Lorena Flores, Ana Vanessa Alonso and Maria Villalba for echocardiography technical assistance, Raquel Baeza and Mercedes de la Cueva for animal housing and handling, Antonio de Molina for histopathological analysis and to CNIC Microscopy and Dynamic Imaging (ICTS-ReDib, cofounded by MCIN/AEI /10.13039/50110001103), Genomics and Histology Core Facilities for technical assistance. B.E, I.M-M and T.A-G performed experiments, analyzed data and helped to write the manuscript. B.P, S.M-T and C.D-D performed experiments. J.R-C performed and analyzed magnetic resonance imaging and coronary resin cast filling by PET-CT. L.J.J.B performed and analyzed echocardiographic studies and coronary diameter analysis and provided essential cardiological feedback for the project. M.C.C provided critical discussion of experiments and the manuscript. K.T provided Kawasaki human tissue and critical discussion of the results. S.M-P defined the concept; planned, performed, and supervised experiments; analyzed data; wrote the manuscript; composed figures together with I.M-M and T.A-G and obtained funding to support the study.

