## Supplemental Data for "Activation of HIF2 leads to vascular remodeling and inflammation, coronary thrombosis and arterial dilation, recapitulating cardiac involvement of Kawasaki disease"

SUPPLEMENTARY FIGURE LEGENDS

FIGURE S1

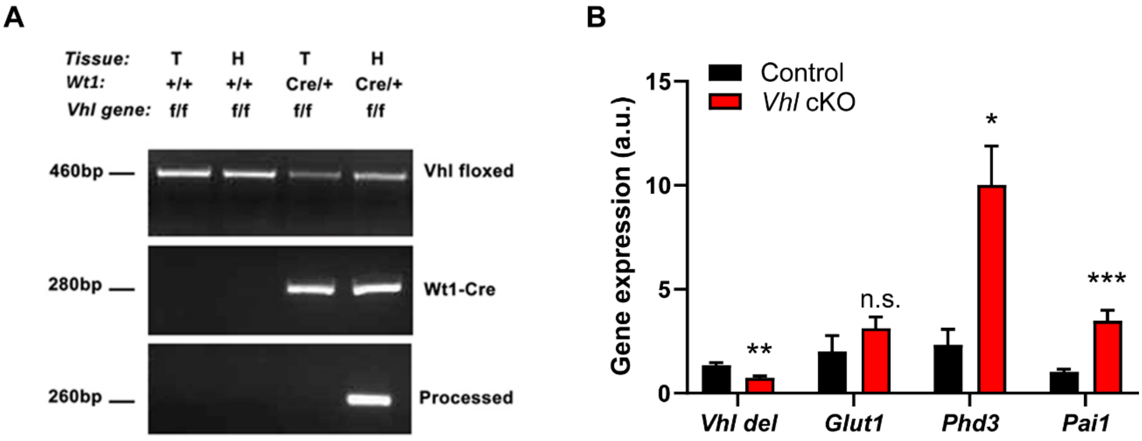


**Figure S1: Efficiency of Vhl locus deletion in Vhl/Wt1Cre mice. A).** Agarose electrophoresis showing PCR products from tail (T) and heart (H) tissue of adult control embryos ($Vhl^{f/f}/Wt1^{+/+}$, lanes 1 and 2) and *Vhl* cKO mutants ($Vhl^{f/f}/Wt1^{Cre/+}$, lanes 3 and 4). Top gel: floxed (460 bp) allele of the *Vhl* gene. Middle gel: Cre (280 bp) allele of the *Wt1* gene. Bottom gel: processed *Vhl* allele after Cre-mediated recombination (260 bp). **B)** Gene expression analysis by qRT-PCR of the levels of the floxed exon of *Vhl* and the HIF target genes *Glut1, Phd3* and *Pai-1* from control (black bars) and *Vhl* cKO (red bars) adult hearts. Bars represent mean±SEM. *Vhl del:* control (n=3), *Vhl* cKO (n=4); *Glut1:* control (n=7), *Vhl* cKO (n=4); *Pghd3: control* (n=7), *Vhl* cKO (n=5); *Pai1*: control (n=6), *Vhl* cKO (n=5). n.s: non-significant; *P≤0.05; **p≤0.01; ***P≤0.001. Statistical significance was tested by Student´s t test.

**FIGURE S2**

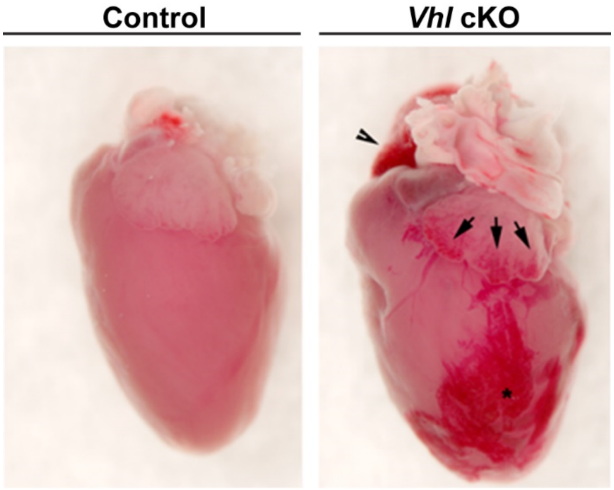


**Figure S2: *Vhl*-deficient mice show cardiomegaly and visible vascular lesions in a macroscopic analysis.** Representative lateral whole mount views of hearts from control and *Vhl* cKO mice. In the *Vhl* cKO arrowhead indicates right atria dilation, arrows point to epicardial vascular lesions in the atria and asterisk show vascular defects in the ventricle.

**FIGURE S3**

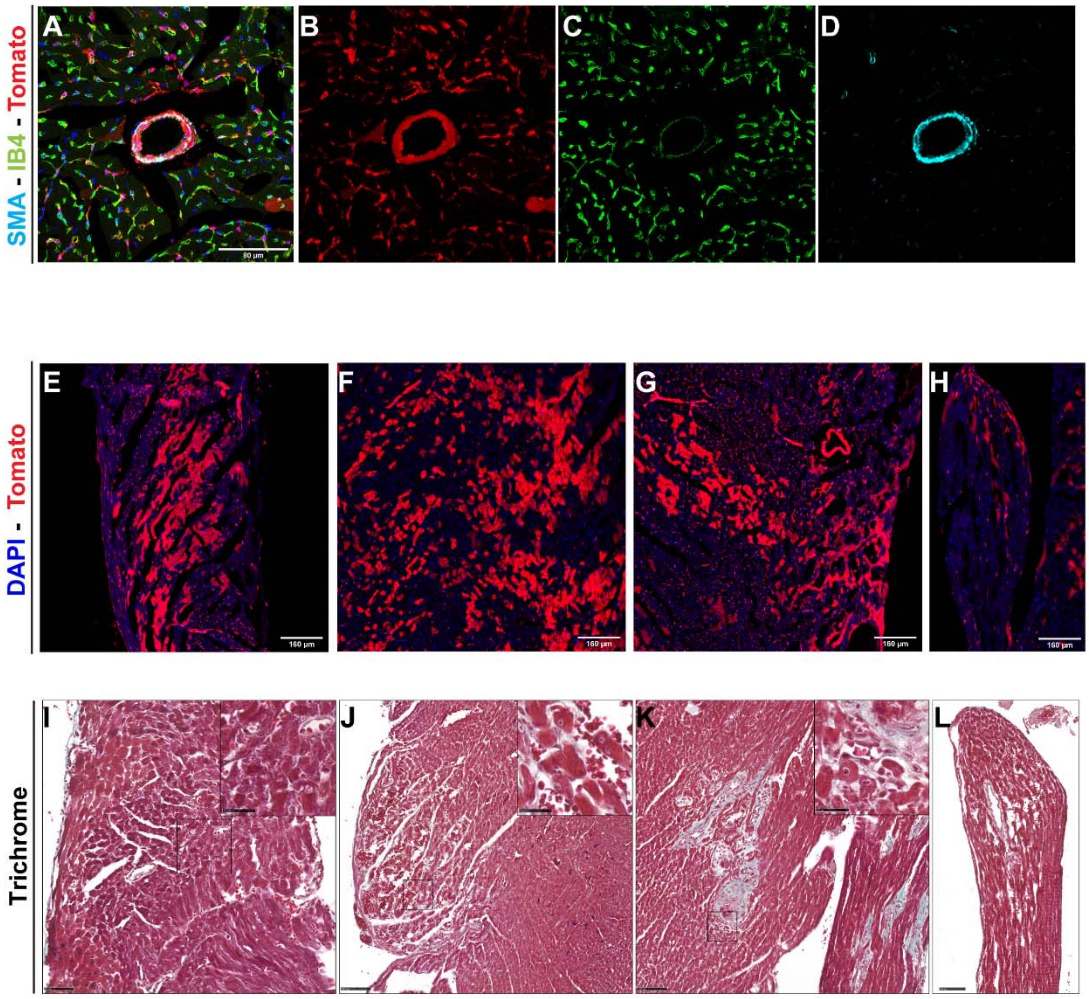


**Figure S3: Wt1 cardiac lineage tracing and defects associated with *Vhl* deletion in Wt1 positive cells**. **A-D)** Immunofluorescence of vascular markers in a cardiac section from Rosa26-tdTomato/Wt1-Cre reporter mice to show contribution to coronary vessels and microvasculature. **B**) Tomato channel (red) reporting cells derived from Wt1+ progenitors or where Wt1 is expressed. **C**). Isolectin B4 (IB4) staining (green) labelling ECs. **D**). Smooth muscle actin (SMA) staining (cyan) labelling the medial layer of arterial coronaries and to lesser extent some pericytes of the microvasculature. **A**). Composite of Tomato, Dapi, IB4 and SMA channels showing positive contribution of the Wt1 lineage to coronary ECs and VSMCs as well as to ventricular microvasculature. Scale bar 80µm. **E-H)** Representative cardiac sections from the Rosa26-tdTomato/Wt1-Cre reporter mice showing Tomato signal in patches of CMs (red) in the right ventricle (**E**), interventricular septum (**F**), left ventricle (**G**) and papillary muscles (**H**). Scale bars 160 µm. These patches correspond to the zones with vascular lesions in the *Vhl* cKO heart (**I-L**). **I-L)** Representative 4 weeks old *Vhl* cKO mice histological sections analyzed by Masson´s trichrome staining showing the initiation of fibrosis and tissue damage in patches of CMs in the right ventricle (**I**), interventricular septum (**J**), left ventricle (**K**) and papillary muscles (**L**) reminiscent of the Wt1 contribution (**E-H**). Scale bars are 50µm (main panels) and 25µm (insets).

**FIGURE S4**

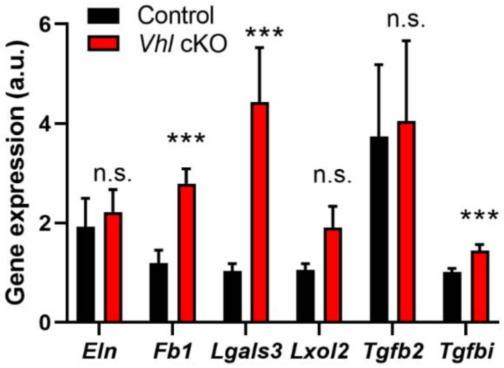


**Figure S4: Gene expression analysis of fibrosis and extracellular matrix remodeling markers.** Relative gene expression analysis determined by RT-qPCR of different target genes involved in fibrosis, vascular remodeling and myofibroblast activation in 8 weeks old hearts. Bars represent mean±SEM*. Eln:* control (n=9), *Vhl* cKO (n=7); *Fb1:* control (n=8), *Vhl* cKO (n=6); *Lgals3: control* (n=6), *Vhl* cKO (n=6); *Lxol2*: control (n=6), *Vhl* cKO (n=6); *Tgfb2* control (n=12), *Vhl* cKO (n=10); *Tgfbi*: control (n=9), *Vhl* cKO (n=9). n.s: non-significant; ***P≤0.001. Statistical significance was tested by Student´s t test.

**FIGURE S5**

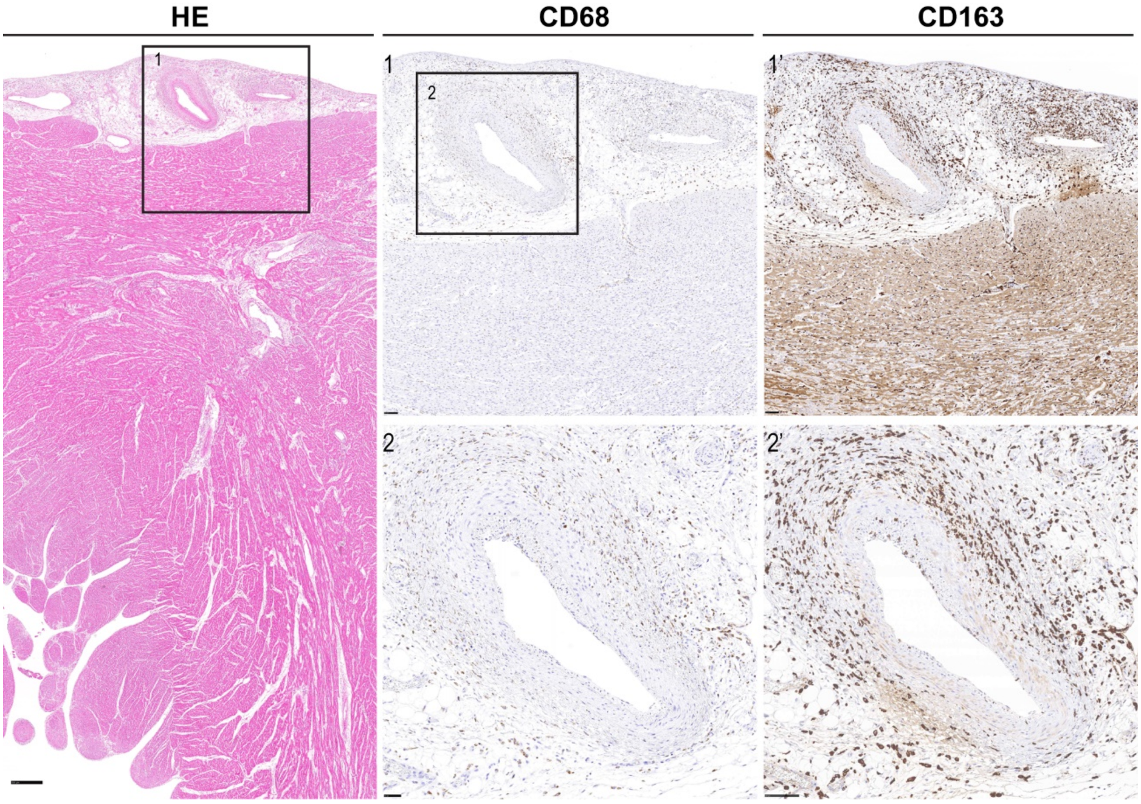


**Figure S5. Perivascular inflammation by infiltration of macrophages in a 7 months old male case of Kawasaki disease.** Left panel correspond to Histological analysis by hematoxylin and eosin staining of a cardiac section from a necropsy of a 7 months old male case of Kawasaki disease with dilated and remodeled epicardial coronary arteries (Left panel. Scale bar 250μm). Inset 1 contains the epicardial zone of the magnifications of the top panels stained for CD68 (1) and CD163 (1´). Inset 2 contains the remodeled artery shown in inset 1 stained with CD68 (2) and CD163 (2´). Scale bars 50μm (2) and 100μm (1, 1´, 2´).
